# Differential Effects of Prostaglandin E_2_ Production and Signaling through the Prostaglandin EP3 Receptor on Human Beta-cell Compensation

**DOI:** 10.1101/670000

**Authors:** Nathan A. Truchan, Harpreet K. Sandhu, Rachel J. Fenske, Renee Buchanan, Jackson Moeller, Austin Reuter, Jeff Harrington, Michelle E. Kimple

## Abstract

**Objective:** Signaling through Prostaglandin E3 Receptor (EP3), a G protein-coupled receptor for E series prostaglandins such as prostaglandin E_2_ (PGE_2_), has been linked to the beta-cell dysfunction and loss of beta-cell mass in type 2 diabetes (T2D). In the beta-cell, EP3 is specifically coupled to the unique cAMP-inhibitory G protein, G_z_. Divergent effects of EP3 agonists and antagonists or Gα_z_ loss on beta-cell function, replication, and survival depending on whether islets are isolated from mice or humans in the lean and healthy, type 1 diabetic, or T2D state suggest a divergence in biological effects downstream of EP3/Gα_z_ dependent on the physiological milieu in which the islets reside.

**Methods:** We determined the expression of a number of genes in the EP3/Gα_z_ signaling pathway; PGE_2_ production pathway; and the beta-cell metabolic, proliferative, and survival responses to insulin resistance and its corresponding metabolic and inflammatory derangements in a panel of 80 islet preparations from non-diabetic human organ donors spanning a BMI range of approximately 20-45. In a subset of islet preparations, we also performed glucose-stimulated insulin secretion assays with and without the addition of an EP3 agonist, L798,106, and a glucagon-like peptide 1 receptor agonist, exendin-4, allowing us to compare the gene expression profile of each islet preparation with its (1) total islet insulin content (2), functional responses to glucose and incretin hormones, and (3) intrinsic influence of endogenous EP3 signaling in regulating these functional responses. We also transduced two independent islet preparations from three human organ donors with adenoviruses encoding human Gα_z_ or a GFP control in order to determine the impact of Gα_z_ hyperactivity (a mimic of the T2D state) on human islet insulin content and functional response to glucose.

**Results:** In contrast to results from islets isolated from T2D mice and human organ donors, where PGE_2_-mediated EP3 signaling actively contributes to beta-cell dysfunction, PGE_2_ production and EP3 expression appeared positively associated with various measurements of functional beta-cell compensation. While Gα_z_ mRNA expression was negatively associated with islet insulin content, that of each of the Gα_z_-sensitive adenylate cyclase (AC) isoforms were positively associated with BMI and cyclin A1 mRNA expression, suggesting increased expression of AC1, AC5, and AC6 is a compensatory mechanism to augment beta-cell mass. Human islets over-expressing Gα_z_ via adenoviral transduction had reduced islet insulin content and secretion of insulin in response to stimulatory glucose as a percent of content, consistent with the effects of hyperactivation of Gα_z_ by PGE2/EP3 signaling observed in islets exposed to the T2D physiological milieu.

**Conclusions:** Our work sheds light on critical mechanisms in the human beta-cell compensatory response, before the progression to frank T2D.

## 1. Introduction

Fundamentally, Type 2 diabetes mellitus (T2D) results because of a failure of the insulin-secreting pancreatic beta-cells to compensate for peripheral insulin resistance. Obesity is the most common co-morbidity found in individuals with insulin resistance. While there exists debate about whether one condition precedes the other, the physiological changes that often occur in the obese, insulin-resistant state (e.g., systemic inflammation, dyslipidemia, hyperinsulinemia, mild hyperglycemia) simultaneously induce beta-cell stress and increase beta-cell workload, forcing the beta-cell to initiate a compensatory response in order to function, replicate, and survive. Whether the beta-cell can initiate and continue this adaptive program is the key determinant of the progression to T2D.

Cyclic AMP (cAMP) is a well-characterized potentiator of glucose-stimulated insulin secretion (GSIS) and promotes a number of proliferative and survival pathways in the beta-cell. Numerous changes in beta-cell cAMP homeostasis occur in a highly-compensating beta-cell, all with the central theme of increasing cAMP production, decreasing cAMP degradation, or promoting signaling through downstream effectors!!refs. Previous work from our laboratory revealed islets from Black and Tan Brachyury (BTBR) mice homozygous for the *Leptin*^*Ob*^ mutation (BTBR-Ob), a strong genetic model of T2D, have a decreased ability to up-regulate cAMP production[1]. BTBR-Ob islets are also less responsive to agents that act through the cAMP-stimulatory glucagon-like peptide 1 receptor (GLP1R), suggesting a tonic brake on cAMP production that interferes with their ability to respond appropriately to glucose [1]. In contrast to GLP1R, Prostaglandin EP3 receptor (EP3), when constitutively active or bound by E-series prostaglandins such as prostaglandin E_2_ (PGE_2_: the most abundant natural ligand for EP3), negatively regulates cAMP production. Islets isolated from T2D mice and human organ donors express more EP3 and produce more PGE_2_ than islets isolated from non-diabetic controls [1], and treating these islets with an EP3 receptor antagonist, L798,106, potentiates GSIS and restores their insulin secretory response to glucose and GLP1R agonists [1].

In beta-cells, EP3 is coupled specifically to the unique inhibitory G protein, G_z_ [2, 3]. Islets from mice lacking the catalytic alpha-subunit of G_z_ (Gα_z_) constitutively produce more cAMP and secrete more insulin in response to glucose [2, 4-6]. Gα_z_-null mice are resistant to diabetes in a number of mouse models of the disease [2, 4, 5]. Gα_z_-null C57BL/6N mice are fully resistant to developing fasting hyperglycemia and glucose intolerance after a high-fat diet (HFD) feeding regimen—a model of pre-diabetes—due primarily to a synergistic effect of HFD feeding and the Gα_z_-null mutation on beta-cell proliferation [2]. While the GSIS response as a percent of islet insulin content is not as dysfunctional as that of wild-type HFD-fed mice, Gα_z_-null islets from HFD-fed mice secrete more insulin primarily because they are larger. This is in contrast to islets from lean, healthy, Gα_z_-null mice or in models of type 1 diabetes (T1D), which secrete more insulin as a percent of content. Gα_z_ loss has no impact on food intake, body weight, insulin sensitivity, or the increase in plasma PGE_2_ metabolite levels induced by HFD feeding, and Gα_z_ tissue distribution is quite limited, the protected phenotype of HFD-fed Gα_z_-null mice is best explained by a beta-cell-centric model.

Our previous studies using mouse models and pancreatic islets from T2D human organ donors, along with an extensive body of work from other investigators, suggest that both PGE_2_ production and EP3 expression are up-regulated by high glucose, pro-inflammatory cytokines, and/or free fatty acids, negatively influencing the islet function and mass. As GPCRs form the largest class of druggable targets in the human genome, we have previously proposed the EP3/Gα_z_ interaction as a novel preventative or therapeutic target for T2D. Yet, a comprehensive analysis of the PGE_2_ production and EP3/Gα_z_ signaling pathways and their correlation with beta-cell function and mass has never been completed in pancreatic islets of non-diabetic human donors; thus, whether targeting the EP3/Gα_z_ interaction would prevent the progression from insulin resistance to T2D remains unclear.

In this work, we determined the mRNA expression of proteins involved in systemic inflammation, beta-cell compensation, PGE_2_ production, and EP3/Gα_z_ signaling in two panels of pancreatic islets isolated from human organ donors (comprising 80 discrete individuals) and, in a sub-set of samples, correlated BMI and gene expression with islet insulin content and different measurements of glucose-stimulated and incretin-potentiated GSIS. We also transduced isolated human islets with an adenovirus expressing human Gα_z_ as a mimic of the chronic up-regulation of EP3/Gα_z_ signaling in the T2D state, quantifying the impact on islet insulin content and GSIS. Our results shed new light on the role of PGE_2_ production and Gα_z_ signaling in the islet compensatory response to the pathophysiological conditions of obesity and insulin resistance and suggest a protective role of PGE2 production and EP3/Gα_z_ signaling the beta-cell highly compensating for the peripheral insulin resistance, metabolic derangements, and inflammation often found in the obese state.

## 2. Materials & Methods

### 2.1. Materials and Reagents

Sodium chloride (S9888), potassium chloride (P3911), magnesium sulfate heptahydrate (M9397), potassium phosphate monobasic (P0662), sodium bicarbonate (S6014), HEPES (H3375), calcium chloride dehydrate (C3881), exendin-4 (E7144) and RIA-grade bovine serum albumin (A7888) were purchased from Sigma Aldrich (St. Louis, MO, USA). Anti-insulin antibodies (Insulin + Proinsulin Antibody, 10R-I136a; Insulin + Proinsulin Antibody, biotinylated, 61E-I136bBT) were from Fitzgerald Industries (Acton, MA, USA). The 10 ng/ml insulin standard (8013-K) and assay buffer (AB-PHK) were from Millipore. RPMI 1640 medium (11879–020: no glucose), penicillin/streptomycin (15070–063), and fetal bovine serum (12306C: qualified, heat inactivated, USDA-approved regions) were from Life Technologies (Carlsbad, CA, USA). Dextrose (D14–500) was from Fisher Scientific (Waltham, MA). The RNeasy Mini Kit and RNase-free DNase set were from Qiagen. High-Capacity cDNA Reverse Transcription Kit was from Applied Biosystems. FastStart Universal SYBR Green Master mix was from Roche (Indianapolis, IN).

### 2.2. Human islet preparations

Cultured human islets were obtained from the Integrated Islet Distribution Program (IIDP) and flash-frozen islet samples from BetaPro according to an approved IRB exemption protocol (UW 2012–0865). The islet preparations were collected in two sets of 40 preparations each: Set 1 was collected between October, 2010 and February, 2012. Islets in this set were only used for gene expression analyses. Set 2 was collected between March, 2013 and May 2015, and were used for both gene expression and insulin secretion assays (The unique identifiers for each islet preparation, along with age, sex, BMI, HbA1c, origin, islet isolation center, and assay the islets were used for are listed in Supplementary Tables 1 and 2). Most of the islet samples for quantitative PCR analysis were hand-picked and pelleted on the day of arrival and flash-frozen prior to RNA isolation. In other samples for qPCR and prior to all *ex vivo* insulin secretion assays, human islets were cultured for at 24 h in human islet medium (RPMI 1640 medium containing 8.4 mM glucose, supplemented with 10% heat-inactivated FBS and 1% penicillin/streptomycin).

### 2.3. Ex vivo islet insulin secretion assays for BMI panel

Isolated human islets used in insulin secretion assays were obtained from the Integrated Islet Distribution Program (IIDP) between 2013 and 2015. The day before assay, islets were hand-picked into 96-well V-bottom tissue culture plates and incubated overnight in RPMI growth medium to adhere islets to the bottom of the plate and single-islet GSIS assays performed with and without the addition of the indicated compounds to the stimulation medium, as previously described [7]. The secretion medium was removed via a multi-channel pipette, and islets were lysed in RLT buffer, resuspended, and secreted insulin and total islet insulin concentration quantified with in-house-generated insulin sandwich ELISA as described previously [7]. In general, secretion medium was diluted 1:20 and content medium diluted 1:200 in order for readings in the linear range of the assay.

### 2.4. Quantitative PCR assays

150-200 islets from each human islet preparation were washed with PBS and used to generate RNA samples via Qiagen RNeasy Mini Kit according to the manufacturer’s protocol. Copy DNA (cDNA) was generated and relative qPCR performed via SYBR Green assay using primers validated to provide linear results upon increasing concentrations of cDNA template, as previously described [5].

### 2.5. Human islet adenoviral transduction and insulin secretion assays

A bicistronic adenovirus expressing GFP and human Gα_z_ was created by subcloning the Gα_z_ coding sequence from a pcDNA 3.1 vector (Bloomsburg University cDNA Resource Center) into the VQpacAd-CMVK-NpA adenoviral entry vector (Viraquest Inc, North Liberty, IA). The entry vector was sent to Viraquest for recombination with a packaging vector bicistronically encoding an enhanced GFP protein (eGFP) and virus amplified and purified. The GFP control virus (VQAd-CMV-eGFP) was purchased from Viraquest. Viraquest viruses are confirmed to be viral E1A-protein free.

On the day of receipt, 1000 islets and shipping media were transferred to a 50 ml conical tube and centrifuged at 800 × g for 2 min. Shipping media was aspirated and the islet pellet was resuspended in 20 ml of fresh islet culture and allowed to recover overnight in 10 cm petri dishes. Islets were pooled into a single 2.5 cm petri dish and hand-picked into a 15 ml conical tube containing 1 ml of a mild Accutase solution and incubated for 30 sec at 37 °C. The Accutase reaction was immediately blocked by addition of islet culture medium, and 300 islets were picked into triplicate petri dishes containing medium with 5 × 10^9^ viral particles/ml (Experiment 1) or 5 × 10^12^ viral particles/ml (Experiments 2 & 3) and cultured for 48 h prior to assay. Two dishes of islets per preparation was used for GSIS assays and one was used to generate cDNA for qPCR. GFP expression was confirmed by fluorescence microscopy and Gα_z_ mRNA expression by qPCR.

On the day of assay, 10 islets per dish were picked into each well of a 12-well multi-well plate, in which each well contained 1 ml modified Krebs Ringer Bicarbonate Buffer (KRBB) containing 0.5% BSA and 1.7 mM glucose. Islets were incubated at 37°C and 5% CO_2_ for 45 min, then transferred into fresh 1.7 mM glucose KRBB and incubated for an additional 45 min. Finally, islets were picked into KRBB containing 1.7 mM glucose or 16.7 mM glucose and incubated again for 45 min. At the end of the assay, islets were picked into a conical tube containing 1 ml PBS, pelleted by pulsing in a microfuge, the PBS removed, and islets lysed in 1 ml of a detergent-based lysis buffer containing 20 mM Tris-HCl, pH 7.5, 150 mM NaCl, 1 mM EDTA, and 1% Triton-X. The stimulation buffer was saved for quantification of secreted insulin. Each treatment condition was performed in duplicate using two independent pools of islets from each donor, providing 6 total biological replicates. The insulin ELISA was performed as described above for the BMI study. GSIS% in 1.7 mM glucose, GSIS% in 16.7 mM glucose, and stimulation index were used in comparative analyses. The unique identifiers for the three islet preparations used in adenoviral infection experiments, along with age, sex, BMI, HbA1c, origin, islet isolation center, are listed in Supplementary Table 3.

### 2.6. Statistical analyses

Data are expressed as mean ± standard error of the mean (SEM) unless otherwise noted. Data were analyzed as appropriate for the experimental design and as indicated in the text and figure legends. In most cases, P-values are reported independent of a statement regarding statistical significance. All of the results of our statistical analyses can be found in table format, and any relationships specifically highlighted in the text as being positively or negatively correlated are also provided in figure format. Statistical analyses were performed with GraphPad Prism version 8 (GraphPad Software, San Diego, CA).

## 3. Results

### 3.1. Islet preparations used in gene expression and insulin secretion assays

#### 3.1.1. Donor Demographics

The gene expression analyses performed in this work used islets isolated from a panel of 80 organ donors spanning a BMI range of approximately 19-45 (See Supplementary Tables 1 and 2). The BMI panel consists of two separate sets of samples: Set 1 was collected from 2011 to 2013 and includes 40 individuals with a BMI of 19-41.3, while Set 2 was collected from 2013-2015 and consists of 40 individuals with a BMI of 22.8-44.7. The two islet sets were well-matched for donor age BMI (Table 1: P=0.67 and 0.47, respectively, by two-tailed t-test), as well as gender distribution (Set 1: N=15 female donors and N=25 male donors; Set 2: N=13 female donors and 27 male donors). Thirteen of 80 islet preparations were from lean donors (BMI < 25), with no donors being underweight (BMI < 18.5); 21 were from overweight donors (BMI 25-29.9); and 46 were from obese donors, with only 6 classified as high-risk (morbid) obesity (BMI ≥ 40). This BMI distribution suggests our panel as a good representation of the normal physiological changes islets are exposed to during the progression from lean to overweight to obesity in both sexes.

**Table 1:** Comparisons between the age and sex distributions of the two sets of islet preparations used in this work

|  | **Set 1 (N=40)** |  |  | **Set 2 (N=40)** |  |  | **P-value** |
| --- | --- | --- | --- | --- | --- | --- | --- |
|  | **Range** | **Mean** | **SD** | **Range** | **Mean** | **SD** |  |
| **Age (yrs)** | 16-65 | 41.4 | 13.2 | 19-63 | 42.6 | 11.6 | 0.67 |
| **BMI (kg/m<sup>2</sup>)** | 19.0-41.3 | 30.4 | 6.0 | 22.8-44.7 | 31.5 | 5.5 | 0.41 |

#### 3.1.2 Genes probed in Islet cDNA Set 1 and Set 2

For qPCR assays, not all genes were tested in both sets, and not every gene was tested in each cDNA sample. Genes unique to Set 1 include adenylate cyclase 1 (*ADCY1), ADCY5, ADCY6*, glucokinase (*GCK*), pyruvate kinase M1/M2 (*PKM*), and cyclin A1 (*CCNA1*). Genes unique to Set 2 include prostaglandin E synthase 3 (*PTGES3*), Gα_z_ (*GNAZ*), and interleukin 6 (*IL6*). Prostaglandin EP3 receptor (*PTGER3*), COX-1 (*PTGS1*) (officially known as prostaglandin-endoperoxidase synthase 1: PTGS1), COX-2 (*PTGS2*) (officially known as PGTS2), *PTGES*, and *PTGES2* were probed in both sets. A table of qPCR primer sequences can be found in Table 2.

**Table 2:**
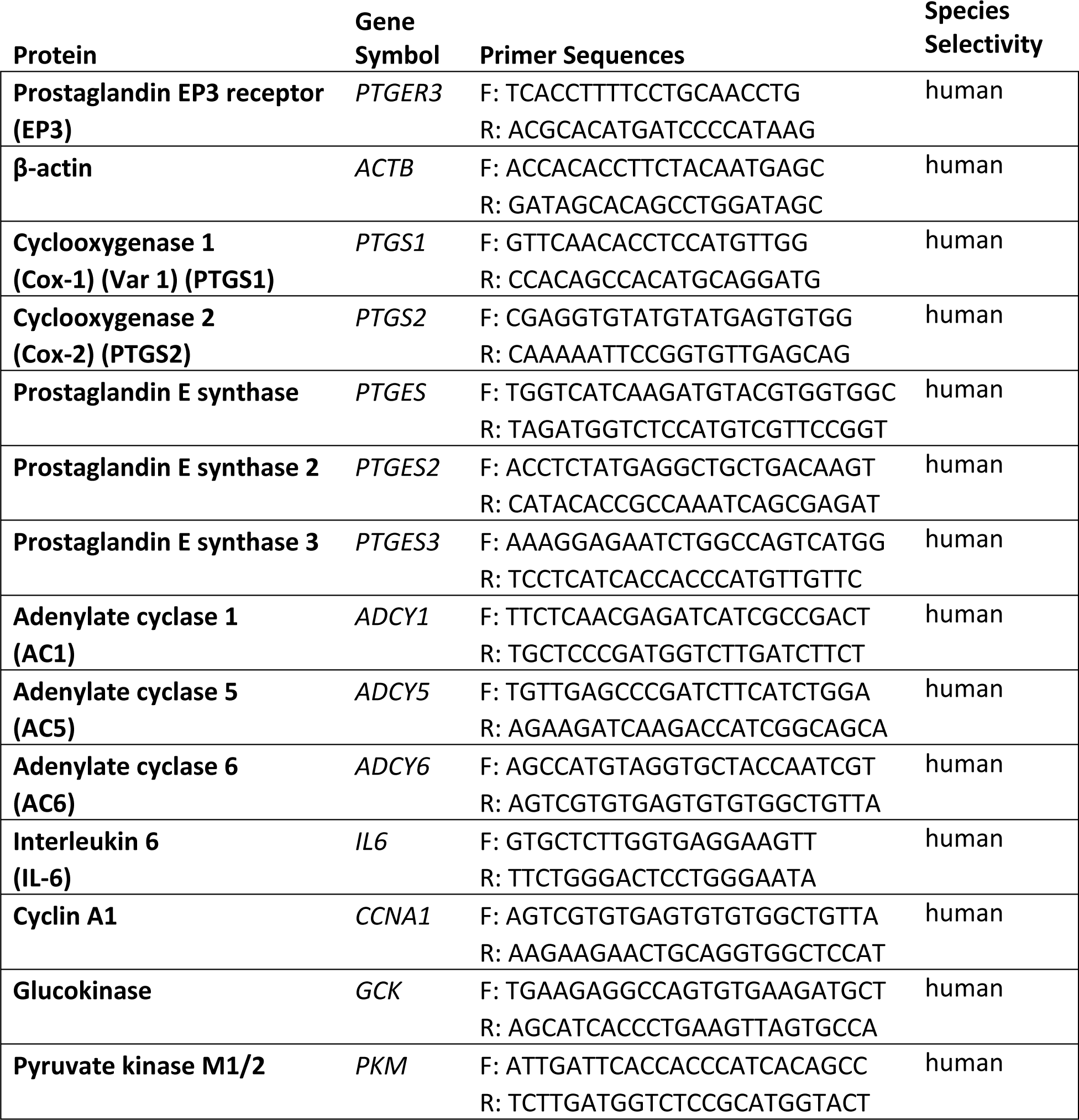
Quantitative PCR primer sequences

#### 3.1.3 Preparations used for insulin secretion assays

Functional assays in response to 1.7 mM glucose and 16.7 mM glucose, with and without the addition of the GLP1R agonist, exendin-4 (10 nM) or the EP3 antagonist, L798,106 (10 µM), were performed on 22 islet preparations from Set 2 of the BMI panel, as indicated in Supplemental Table 2.

### 3.2. Gene/BMI correlations

#### 3.2.1. Overview of analytical methods

We determined the relationship between donor BMI and gene expression using two methods: (1) linear curve-fit analysis and (2) binning samples by donor obesity status (BMI < 30 and BMI ≥ 30; N=17 non-obese and N=23 obese donors in each set) and performing a two-tail T-test. The complete results of this analysis can be found in Supplementary Table 4.

#### 3.2.2. Genes involved in the PGE_2_ production and EP3/Gα_z_ signaling pathways

*PTGER3* expression was not correlated with BMI in either analysis (Supplementary Table 4). Among the genes in the PGE_2_ production pathway tested (*PTGS1, PTGS2, PTGES, PTGES2*, and *PTGES3*), *PTGES2* and *PTGES3* had no apparent relationship with BMI (Supplementary Table 4). *PTGS1* had a weak linear relationship with BMI (p=0.0854, R^2^=0.05397); *PTGS2* had a weak linear relationship with BMI (p=0.043, R^2^=0.0634) and a stronger, positive association with obesity status (p=0.0601); and *PTGES* had a weak linear relationship with BMI (p=0.0733, R^2^=0.04925) (Figure 1A-C). *GNAZ* expression was not correlated with BMI in either analysis (Supplementary Table 4). In contrast, mRNA levels of all three of the Gα_z_-sensitive adenylate cyclase isoforms—*ADCY1, ADCY5*, and *ADCY6*—had positive, linear relationships with donor BMI (*ADCY1*, p=0.0176, R^2^=0.2749; *ADCY5*, p=0.0135, R^2^=0.294; *ADCY6*, p=0.0094; R^2^=0.371); a relationship that, for the most part, also held true when samples were binned by obesity status (p=0.0706, p=0.0380, and p=0.0768, respectively) (Figure 1D-F).

**Figure 1.**
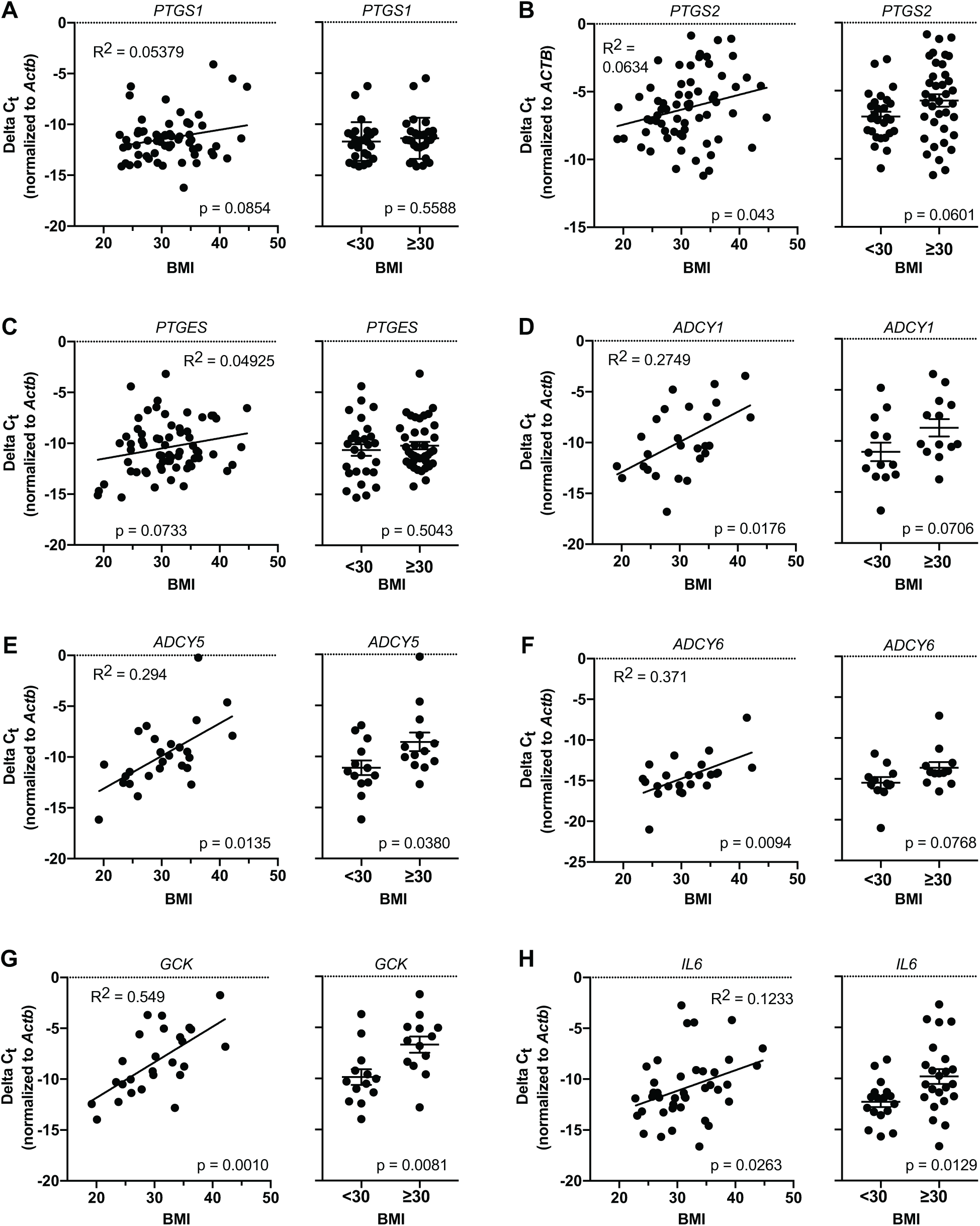
Correlation of donor BMI with islet mRNA expression of proteins involved in the PGE_2_ production pathway, EP3/Gα_z_ signaling pathway, and beta-cell compensatory response. In all panels, relative qPCR was performed with primers specific the indicated genes. Data are represented as ΔC_T_ as normalized to β-actin (*ACTB*), and were subject to linear curve-fit analysis vs. donor BMI (left graphs in each panel) or two-tailed t-test by donor obesity status (right graphs in each panel) in GraphPad Prism version 8 (N = 25-66 for each comparison). For linear curve fit analyses, the goodness-of-fit (R^2^) and p-value for deviation from zero of each of the analyses are indicated. Only relationships with apparent negative or positive correlation are shown in this figure. A-C: PGE_2_ production pathway genes whose expression is correlated with donor BMI. D-F: EP3/Gα_z_ signaling pathway genes whose expression is correlated with donor BMI. G-H: Beta-cell compensation genes whose expression is positively correlated with donor BMI. The full results of this analysis, including genes whose expression is not related to donor BMI, can be found in Supplementary Table 4.

#### 3.2.3. Genes involved in beta-cell compensation

We determined the relative mRNA abundance of proteins important in the adaptive metabolic (*GCK, PKM*), proliferative (*CCNA1*), and survival (*IL6*) response(s) to obesity and insulin resistance as related to donor BMI. Of these, *GCK* and *IL6* had positive linear relationships with BMI (*GCK*, p=0.0010, R^2^=0.549; and *IL6*, p=0.0263; R^2^=0.1233), and the mean expression of *GCK* and *IL6* were elevated in islets from obese donors as compared to lean/overweight (p=0.0081 and p=0.0129, respectively) (Figure 1G,H). *CCNA1* and *PKM* had no apparent relationship with obesity in either analysis (Supplementary Table 4)

### 3.3. Gene/gene correlations

#### 3.3.1. Overview of analytical methods

As with gene/BMI correlations, we determined the relationship between the expression of two genes by linear curve-fit analysis. The complete results of this analysis can be found in Supplementary Table 5.

#### 3.3.2 Genes involved in the EP3/Gα^z^ signaling and PGE_2_ production pathways

We plotted the expression of *PTGER3, ADCY1, ADCY5, ADCY6, PTGS1, PTGS2, PTGES, PTGES2*, PTGES3, and *GNAZ* against each other. All of the genes in the EP3/Gα_z_ signaling pathway had one or more relationships with each other, with *PTGER3* being related to nearly all of the others (*PTGER3* vs. *GNAZ*, p=0.0089, R^2^=0.1751; *PTGER3* vs. *ADCY1*, p=0.0278, R^2^=0.2016; *PTGER3* vs. *ADCY5*, p=0.0106, R^2^=0.2618; *ADCY1* vs. *ADCY5*, p=0.0002, R^2^=0.4617; *ADCY5* vs. *ADCY6*, p=0.0111, R^2^=0.2942) (Figure 2A-E).

**Figure 2.**
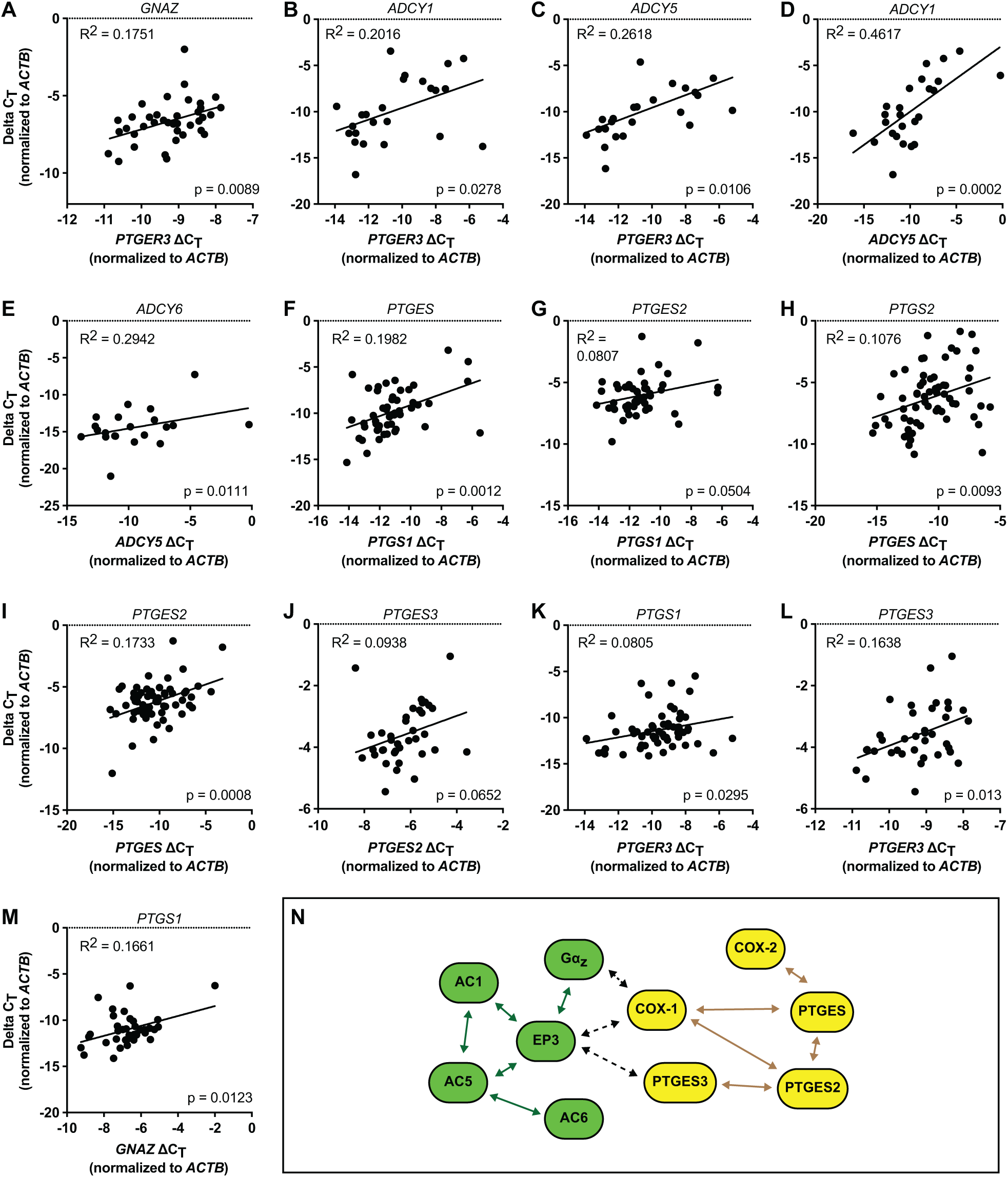
Correlation of mRNA expression of proteins involved in the EP3/Gα_z_ signaling pathway and PGE_2_ production pathway within and across groups. In all panels, relative qPCR was performed with primers specific the indicated genes. Data are represented as ΔC_T_ as normalized to β-actin (*ACTB*), and were subject to linear curve-fit analysis in GraphPad Prism version 8 (N = 13-25 for each comparison). The goodness-of-fit (R^2^) and p-value for deviation from zero of each of the analyses are indicated. Only relationships with apparent negative or positive correlation are shown in this figure. A-E: EP3/Gα_z_ signaling pathway genes as correlated with each other. F-J: PGE_2_ production pathway genes as correlated with each other. K-M: Apparent cross-pathway correlations. N: Summary of the relationships between gene expression of EP3/Gα_z_ signaling pathway proteins (Green) and PGE_2_ production pathway proteins (Yellow) within and across pathways. Within-pathway relationships are indicated by dark green or dark yellow arrows, respectively, and cross-pathway relationships are indicated by dashed black arrows. The full results of this analysis, including genes whose expression is not related to the others, can be found in Supplementary Table 5.

As with the expression genes in the EP3/Gα_z_ signaling pathway, those in PGE_2_ production pathway were also highly correlated with each other (*PTGS1* vs. *PTGES*, p=0.0012, R^2^=0.1982; *PTGS1* vs. *PTGES2*, p=0.0504, R^2^=0.0807; *PTGES vs. PTGS2*, p=0.0093, R^2^=0.1076; *PTGES* vs. *PTGES2*, p=0.0008, R^2^=0.1733; *PTGES2* vs. *PTGES3*, p=0.0652, R^2^=0.0938) (Figure 2F-J).

*PTGER3* and *GNAZ* were the only genes that bridged the two pathway groups, with *PTGER3* being related to *PTGS1* and *PTGES3* (p=0.0295, R^2^=0.0805 and p=0.0103, R^2^=0.1638, respectively), and *GNAZ* being related to *PTGS1* (p=0.0123, R^2^=0.1661) (Figure 2K-M). Figure 2N shows a model of the relationships between the genes in the EP3/Gα_z_ signaling pathway and PGE_2_ production pathway, with two clusters clearly evident. As *ADCY1, ADCY5*, and *ADCY6* were only probed in Set 1, and *GNAZ* and *PTGES3* only in Set 2, it is possible more intra- or inter-cluster connections exist.

#### 3.3.3. Genes involved in beta-cell compensation

*CCNA1, GCK*, and *PKM* expression were all positively correlated with each other (*CCNA1* vs. *GCK*, p=0.0075, R^2^=0.294; *CCNA1* vs. *PKM*, p=0.0847, R^2^=0.1413; *GCK* vs. *PKM*, p=0.0222, R^2^=0.2461) (Table 1). As *IL6* was only probed in Set 2, its relationship with the others was not determined.

Of the genes in the EP3/Gα_z_ signaling pathway and PGE_2_ production pathway, only those involved in EP3/Gα_z_ signaling were correlated with *CCNA1* (*PTGER3*, p=0.04, R^2^=0.1781; *ADCY1*, p=0.0154, R^2^=0.2487; *ADCY5*, p=0.0591, R^2^=0.1594) (Figure 3A-C). *ADCY6* had a weak positive, linear correlation with *CCNA1* (p=0.097, R^2^=0.137) (Supplementary Table 5). *PTGER3, ADCY1, ADCY5*, and *ADCY6* were also the only genes correlated with *GCK*, with all four being positively associated (*PTGER3*, p=0.0003, R^2^=0.4481; *ADCY1*, p=0.0003, R^2^=0.4422; *ADCY5*, p=0.0005, R^2^=0.4165; and *ADCY6*, p=0.0288, R^2^=0.2275) (Figure 3D-G). *PTGER3* was also positively associated with *PKM* (*PTGER3*, p=0.0016, R^2^=0.399) (Figure 3H). In contrast, none of the genes in the PGE_2_ production pathway were correlated with *CCNA1, GCK*, or *PKM* expression (Supplementary Table 5), save a negative correlation between *PTGES* expression and *PKM* (p=0.0226, R^2^=0.3894) (Figure 3I). In contrast, all genes involved in the PGE_2_ production pathway, save *PTGES3*, were positively correlated with *IL6* expression (*PTGS1*, p=0.0057, R^2^=0.1999; *PTGS2*, p=0.0009, R^2^=0.2823; *PTGES*, p=0.0067, R^2^=0.1873; and *PTGES2*, p=0.0395, R^2^=0.1126) (Figure 3K-M). Neither *PTGER3* nor *GNAZ* was correlated with *IL6* expression (Supplementary Table 5). These results suggest PGE_2_ production and signaling through EP3/Gα_z_ both influence beta-cell adaptation to obesity and insulin resistance through at least partially divergent mechanisms. Figure 3N shows a model of the relationships between the genes in the EP3/Gα_z_ signaling pathway and PGE_2_ production pathway with those involved in beta-cell compensation. As with the EP3/Gα_z_ signaling and PGE_2_ production pathway correlations shown in Figure 2N, two clusters were clearly evident. As *CCNA1, GCK, PKM, ADCY1, ADCY5*, and *ADCY6* were only probed in Set 1, and *IL6, GNAZ* and *PTGES3* only in Set 2, it is possible more intra- or inter-cluster connections exist.

**Figure 3.**
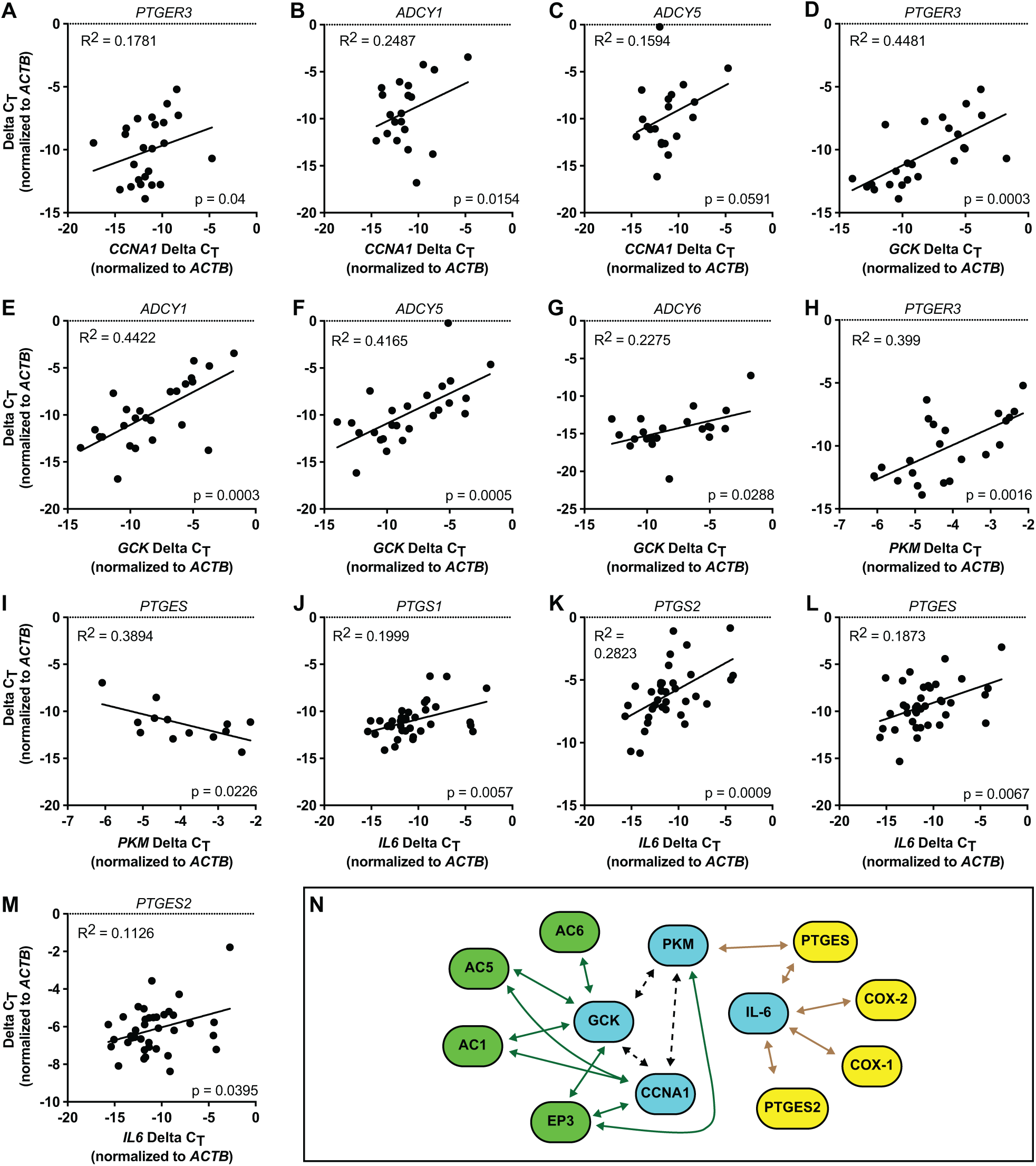
Correlation of mRNA expression of proteins involved in the EP3/Gα_z_ signaling pathway and PGE_2_ production pathway with mRNA expression of markers of beta-cell compensation. In all panels, relative qPCR was performed with primers specific the indicated genes. Data are represented as ΔC_T_ as normalized to β-actin (*ACTB*), and were subject to linear curve-fit analysis in GraphPad Prism version 8 (N = 38-40 for each comparison). The goodness-of-fit (R^2^) and p-value for deviation from zero of each of the analyses are indicated. Only relationships with apparent negative or positive correlation are shown in this figure. A-C: Genes correlated with *CCNA1*. D-F: Genes correlated with *GCK*. H,I: Genes correlated with *PKM*. J-M: Genes correlated with *IL6*. N: Summary of the relationships between gene expression of EP3/Gα_z_ signaling pathway proteins (Green) and PGE_2_ production pathway proteins (Yellow) with markers of beta-cell compensation (Teal). Dashed black arrows indicate the relationships of beta-cell compensation genes with each other. The full results of this analysis, including genes whose expression is not related to the others, can be found in Supplementary Table 5.

### 3.4. Gene/islet insulin content and beta-cell function correlations

#### 3.4.1. Overview of experimental design and analytical methods

For about half of the islet preparations represented in our latter BMI panel (22 of 40), we performed static GSIS assays in 1.7 mM glucose, 16.7 mM glucose, 16.7 mM glucose + 10 nM exendin-4 (Ex4: a GLP1R agonist), 16.7 mM glucose + 10 µM L798,106 (L798: a specific EP3 antagonist), and 16.7 mM glucose + Ex4 + L798. Therefore, we are able to plot BMI or relative expression of the eight genes probed in that panel (*PTGER3, GNAZ, PTGS1, PTGS2, PTGES, PTGES2, PTGES3* and *Il6*) with measurements of beta-cell mass and function: (1) islet insulin content; (2) glucose-stimulated insulin secreted (GSIS); (3) GSIS as a percent of islet insulin content (GSIS%); (4) secretion index (SI), or the ratio of GSIS in 16.7 mM glucose over that in 1.7 mM glucose; (5) incretin response (IR), or the ratio of GSIS in 16.7 mM glucose + Ex4 over than in 16.7 mM glucose alone; (6) L798 SI, or the ratio of GSIS in 16.7 mM glucose + L798 over that in 16.7 mM glucose alone; and (7) L798 IR, or the ratio of GSIS in 16.7 mM glucose + Ex4 + L788,106 over that in 16.7 mM glucose + Ex4. The complete results of this analysis can be found in Supplementary Table 6.

#### 3.4.2 Quality-control measurements and population-based analyses of human islet functional response

Beta-cell function can be compromised in cultured, shipped islets. A Wilcoxon signed-rank test revealed the median SI of the entire islet population (5.90) deviated significantly from 1 (p<0.0001), indicating overall good glucose responsiveness in the majority of islet preparations (Figure 4A). The median IR of all islet preparations (1.27) deviated significantly from 1 (p<0.0001), indicating intact GLP1R signaling in the majority of islet preparations (Figure 4B, left). Because of their opposing effects on cAMP production, agonists of GLP1R and antagonists of EP3 should have additive, positive effects on SI, presuming that agonist-sensitive EP3 isoforms are expressed and are being stimulated by endogenous PGE_2_. The median L798 SI (0.98) was not significantly different from 1 (p=0.6333) (Figure 4B, middle), indicating weak endogenous agonist-dependent EP3 signaling in these islets, as expected based on previous work [1]. The median L798 IR (1.14) did deviate significantly from 1 (p=0.0071) (Figure 4B right), revealing, in the majority of islet preparations, L798,106 treatment ameliorated an (albeit small) inhibitory constraint on cAMP-mediated GLP1R signaling. This result was only revealed by considering the distribution of L798 IR of the entire islet population, as the L798 IR of each islet preparation was not significantly different from baseline IR (data not shown), consistent with our previously-published results using a smaller set of human islet preparations [1].

**Figure 4.**
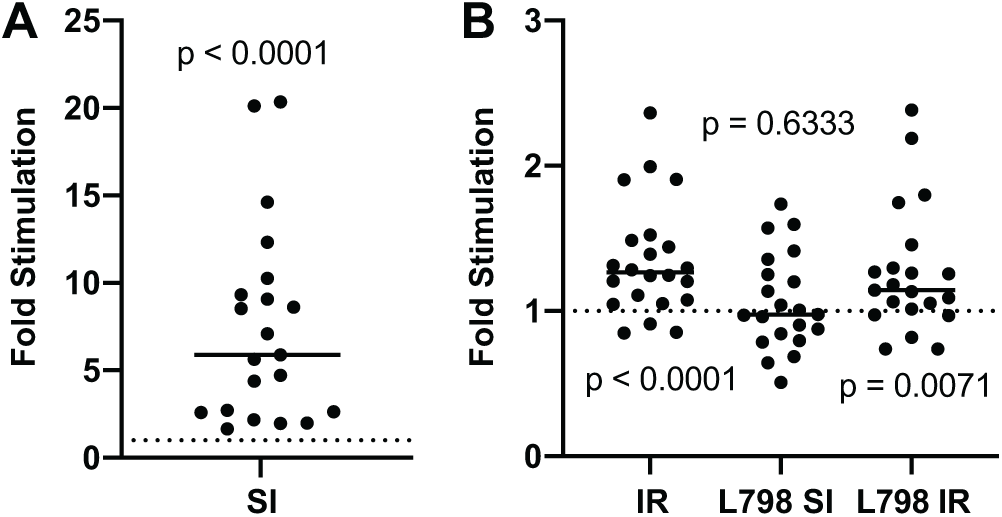
Population analyses of cultured human islet function. A Wilcoxon signed-rank test was performed to determine whether the median response for each condition deviated significantly from a hypothetical value of 1 (e.g., no effect of the treatment on GSIS). A: Stimulation index (SI). B: Incretin response (IR), L798 SI, and L798 IR. The bar in each graph indicates the median value for each measurement. P-values for each analysis are indicated. N = 22 independent islet preparations.

#### 3.4.3 Relationship of donor BMI with islet insulin content and beta-cell function

BMI was positively correlated with islet insulin content (p=0.0042, R^2^=0.3415) (Figure 5A) and GSIS in all treatment conditions (1.7G: p=0.0128, R^2^=0.2718; 16.7G; p=0.0261, R^2^=0.224; 16.7G + Ex4: p=0.0622, R^2^=0.1589; 16.7G + L798: p=0.0107, R^2^=0.2966; and 16.7G + Ex4 + L798: p=0.0055, R^2^=0.3404) (Supplementary Table 6). GSIS% was unaffected by BMI, though (Supplementary Table 6), indicating islets from higher BMI donors secrete more insulin simply because they have more insulin to secrete. A weak, negative correlation of BMI with GSIS% in 1.7 mM glucose (p=0.0839; R^2^=0.1419) suggests possible beta-cell dysfunction in islets from donors with higher BMI (Supplementary Table 6). BMI also had a weak negative correlation with IR (p=0.1219, R^2^ = 0.1154), but not with other measurements of islet responsivity to any other treatment (SI, L798 SI, and L798 IR) (Supplementary Table 6).

**Figure 5.**
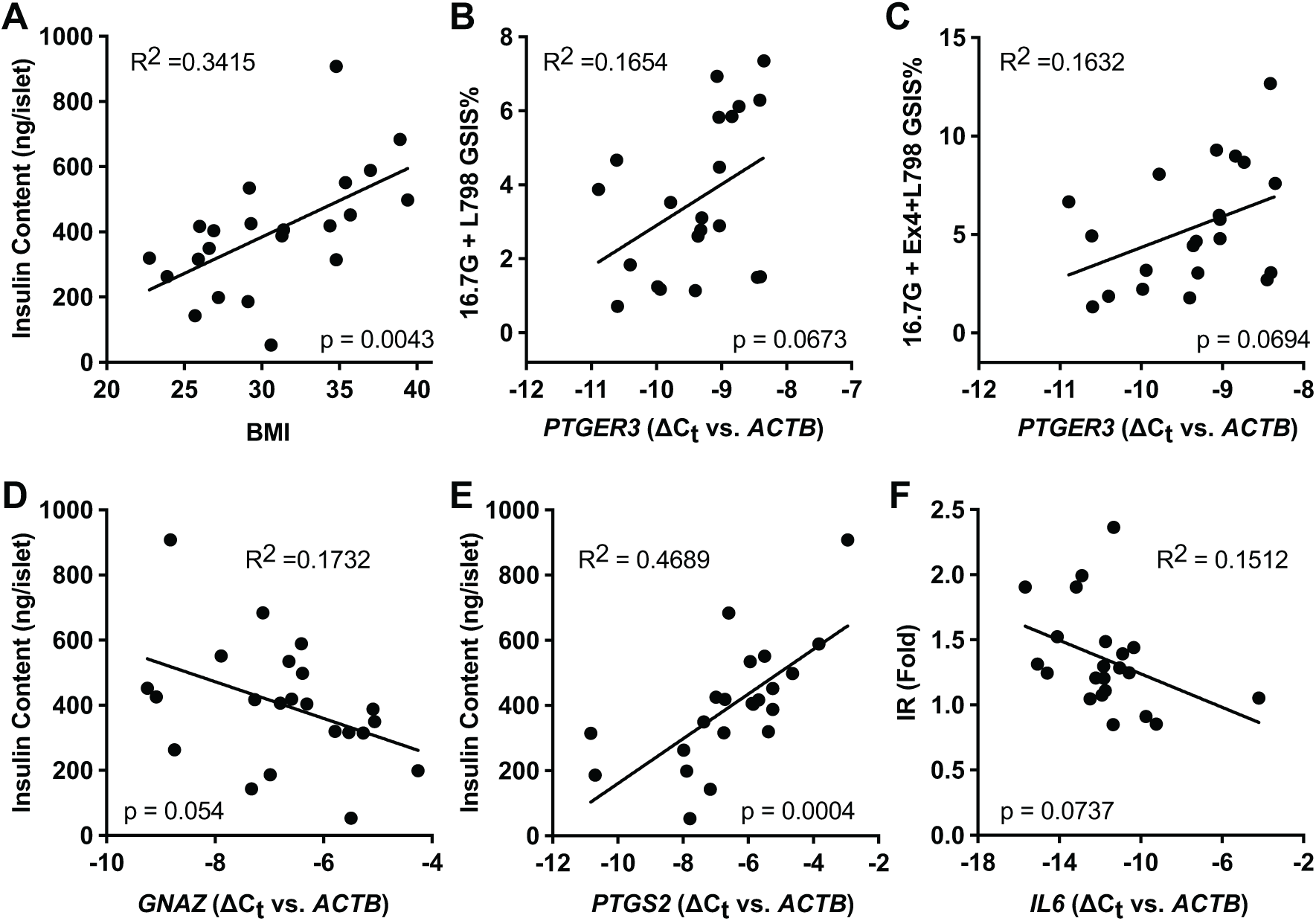
Correlation of BMI or mRNA expression of proteins involved in the EP3/Gα_z_ signaling pathway and PGE_2_ production pathway with islet insulin content and beta-cell function. For each islet preparation in Set 2 subject to GSIS analysis (N=22), donor BMI or expression of *PTGER3, GNAZ, PTGS1, PTGS2, PTGES, PTGES2, PTGES3*, or *IL6*—as determined by relative qPCR with primers specific for the indicated genes and represented as ΔC_T_ as normalized to β-actin (*ACTB*)—were subject to linear curve-fit analysis against total islet insulin content or different measurements of beta-cell function in GraphPad Prism version 8 (N = 13-25 for each comparison). The goodness-of-fit (R^2^) and p-value for deviation from zero of each of the analyses are indicated. Only relationships with apparent positive or negative correlation are shown in this figure. The full results of this analysis can be found in Supplementary Table 6.

#### 3.4.4. Relationship of PTGER3 and GNAZ expression with islet insulin content and beta-cell function

*PTGER3* expression had no relationship with islet insulin content, GSIS, GSIS% in 1.7 mM or 16.7 mM glucose, SI, IR, L798 SI, or L798 IR (Supplementary Table 6). On the other hand, GSIS% in 16.7 mM glucose, with or without the addition or Ex4, was enhanced with L798,106 co-treatment (16.7G + L798: p=0.0673, R^2^=0.1654; 16.7G + Ex4 + L798; p=0.0694, R^2^=0.1632) (Figure 5B,C). These results are consistent with the expected effect of an EP3 antagonist on GSIS% if agonist-sensitive EP3 variants are being stimulated by endogenous PGE_2_ [1].

*GNAZ* expression was negatively correlated with islet insulin content (p=0.054; R^2^=0.1732) (Figure 5D). A negative relationship of *GNAZ* expression on insulin content was reflected in a negative influence on GSIS in 1.7 mM glucose, 16.7 mM glucose, and 16.7 mM glucose + Ex4, although these relationships were overall weak (Supplementary Table 6). *GNAZ* expression did not impact GSIS% in any treatment condition, including with the addition of L798,106, SI, IR, L798 SI, or L798 IR (Supplementary Table 6). In comparison to the results described above for *PTGER3*, these results suggest any reduction of GSIS by increased *GNAZ* expression is primarily driven by decreased insulin content and not by an influence of PGE_2_-mediated EP3 signaling.

#### 3.4.5. PTGS2 expression, and not that of other genes in the PGE_2_ production pathway, is positively correlated with islet insulin content and secretory capacity

Like BMI, *PTGS2* expression was positively correlated with higher islet insulin content (p=0.0004; R^2^=0.4689) (Figure 5E) and total insulin secreted in all conditions (1.7G: p=0.0079, R^2^=0.3037; 16.7G; p=0.006, R^2^=0.3209; 16.7G + Ex4: p=0.015, R^2^=0.2616; 16.7G + L798: p=0.0244, R^2^=0.2394; and 16.7G + Ex4 + L798: p=0.0175, R^2^=0.2629), but not GSIS% (Supplementary Table 6). Unlike BMI, *PTGS2* expression did not influence GSIS% in 1.7 mM glucose or IR (Supplementary Table 6), suggesting *PTGS2* expression alone does not promote beta-cell dysfunction. None of the other genes in the PGE_2_ synthetic pathway (*PTGS1, PTGES, PTGES2*, and *PTGES3*) had any significant impact on islet insulin content, GSIS, GSIS%, SI, IR, L798 SI, or L798 IR (Supplementary Table 6). These results support the key role of COX-2 as the rate-limiting enzyme in PGE_2_ production, with the expression of other enzymes—all being up-regulated by the same inflammatory conditions that induce *PTGS2* expression—being non-rate-limiting.

#### 3.4.6. Exclusion of selection bias in IL6 and PTGS2 GSIS assays

In the full set of islet preparations comprising Set 2, *IL6* expression was positively correlated with BMI (with *PTGS2* more weakly so), and *IL6* and *PTGS2* expression were highly correlated with each other. Yet, only BMI and *PTGS2* expression were positive predictors of islet insulin content: *IL6* expression had no relationship with islet insulin content or any measurements of beta-cell function (Supplementary Table 6). In order to exclude selection bias in these divergent results, we limited our gene expression analyses to only those of the 22 islet preparations used in GSIS assays, and found the same relationships(s), albeit weaker, as when the full set was used (*IL6* vs. BMI linear curve-fit: p=0.1137, R^2^=0.1204; *IL6* by obesity status: p=0.1048; *PTGS2* vs. BMI linear curve-fit: p=0.2028, R^2^=0.07997; *PTGS2* by obesity status: p=0.1626; *IL6* vs. *PTGS2*: p=0.179; R^2^=0.2494) (Figure 6A-E).

**Figure 6.**
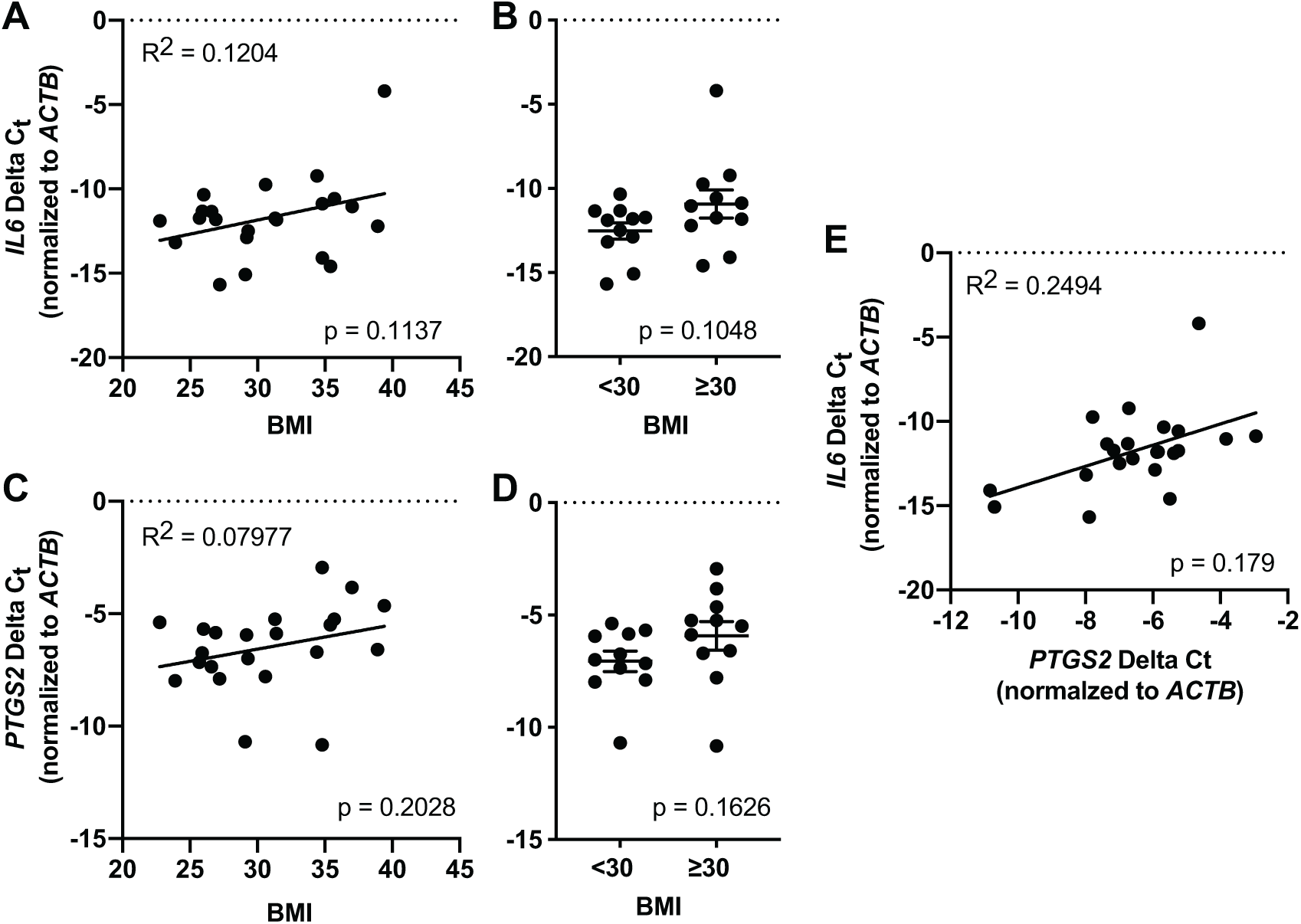
Maintenance of relationship between *IL6* and *PTGS2* mRNA abundance with BMI and each other in the subset of islets used in GSIS assays. A and B: *IL6* abundance vs. donor BMI (A) or obesity status (B). C and D: *PTGS2* abundance vs. donor BMI (C) or obesity status (D). E: *IL6* abundance vs. *PTGS2* abundance. Relative qPCR was performed with primers specific the indicated genes. In all panels, gene expression data are represented as ΔC_T_ normalized to β-actin (*ACTB*). In A, C, and E, linear curve-fit analysis was performed in GraphPad Prism version 8. The goodness-of-fit (R^2^) and p-value for deviation from zero of each of the analyses are indicated. In B and D, the mean relative gene expression in non-obese vs. obese donors was compared by two-way t-test, with the p-value for each analysis indicated. N = 22 for each comparison.

### 3.5. Gα_z_ over-expression in human donor islets and effects on islet insulin content and GSIS

#### 3.5.1. Overview of experimental design and analytical methods

When islets are isolated from T2D human organ donors, tonic activation of EP3 by PGE_2_ blunts the ability of the beta-cells to secrete insulin in response to glucose: a phenotype that can be at least partially ameliorated by the EP3 antagonist, L798,106. To explore the hypothesis that increased Gα_z_ activity in the T2D state is responsible for defects in beta-cell function, we transduced islets from three human donors with an adenovirus bicistronically encoding GFP and human Gα_z_, or a GFP control adenovirus, and cultured islets for two days prior to GSIS assays. Two independent pools of islets from each donor were used, providing 6 biological replicates total. After two days in culture, GFP expression was clearly evident by fluorescence microscopy (Figure 7A), and qPCR confirmed *GNAZ* overexpression (*ACTB* C_t_, ∼21 for all preparations; *GNAZ* C_t_ ∼24, GFP-infected vs. C_t_ ∼15-18, Gα_z_-infected; data not shown). With *GNAZ* C_t_ values outside of the linear range of the assay, the degree of overexpression was not quantified. All islet insulin content and GSIS results were normalized to those of the mean of the GFP control to account for (1) intrinsic differences in total islet insulin content and GSIS% among the three islet preparations and (2) the effect of adenoviral transduction itself. The raw GSIS data can be found in Supplementary Figure 1.

**Figure 7.**
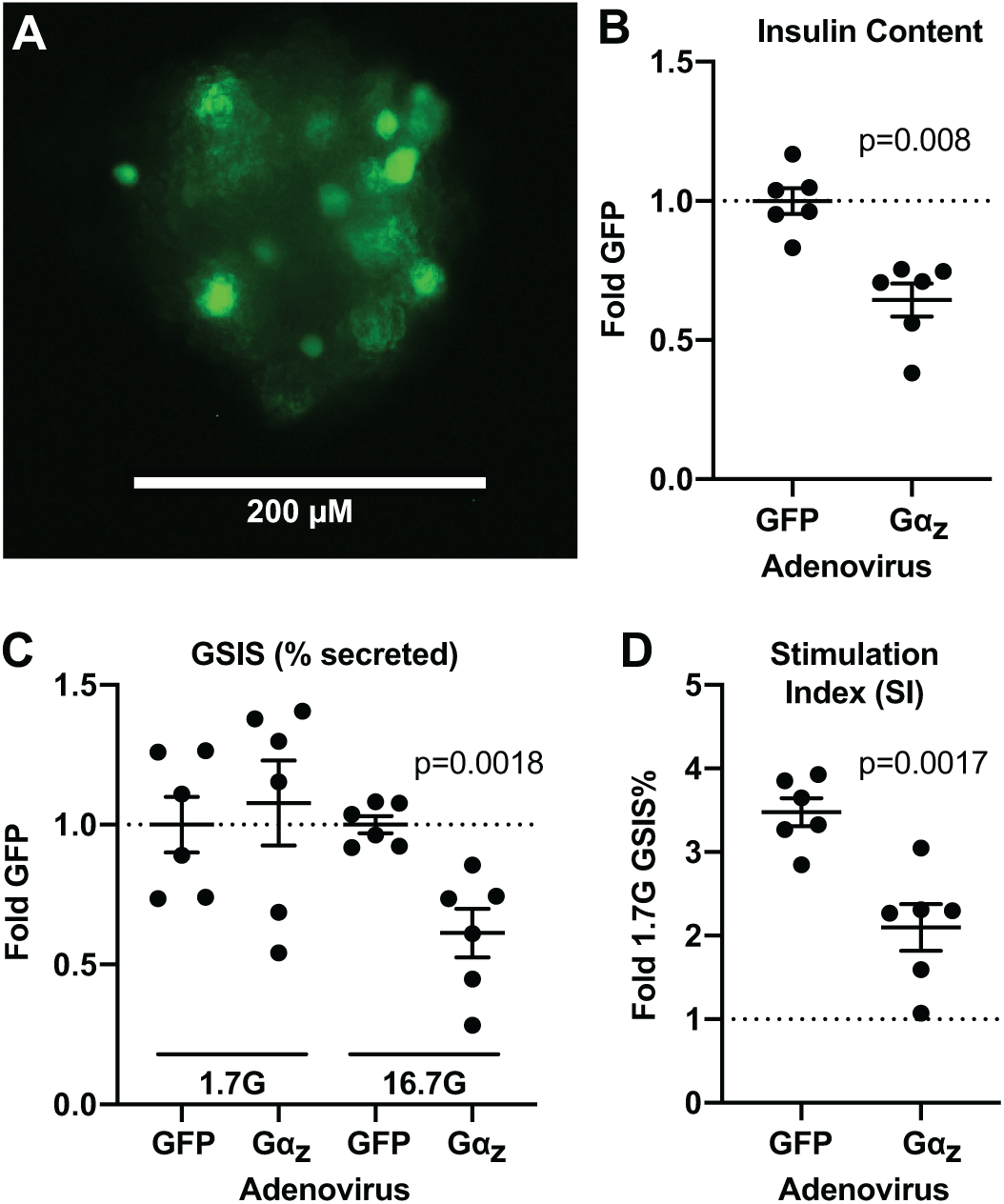
Gα_z_ overexpression reduces both islet insulin content and the percent of insulin secreted in stimulatory glucose in cultured human islets. A: Confirmation of effective adenoviral transduction of cultured human islets by fluorescence microscopy for the eGFP tracer. B: Islet insulin content as normalized to the mean content in GFP-transduced islets for each independent islet preparation. C: GSIS% in 1.7 mM glucose or 16.7 mM glucose as normalized to the mean of GFP-transduced islets for each independent islet preparation. D: Stimulation index (SI) for each independent islet preparation. Data were compared by two-tail t-test in GraphPad Prism v. 8.0, and the p-value indicated on the graph. For all panels, N = 2 individual preparations from 3 separate donors (N = 6 total).

#### 3.5.2. Gα_z_ over-expression in human donor islets mimics the beta-cell dysfunction of T2D

After two days in culture, the mean insulin content was 36% lower in Gα_z_-transduced islets as compared to GFP control (GFP, 0.9999 ± 0.04628, vs. Gα_z_ 0.6438 ± 0.05973; p = 0.008) (Figure 7B). Furthermore, while GSIS% in 1.7 mM glucose was unchanged in Gα_z_-transduced islets as compared to GFP control, GSIS% in 16.7 mM glucose was 39% lower (1.00 ± 0.03063, GFP vs. 0.6127± 0.08711, Gα_z_; p = 0.0018), and the mean stimulation index was 40% lower (3.479 ± 0.1669, GFP vs. 2.009 ± 0.2783, Gα_z_; p = 0.0017) (Figure 7C,D).

## 4. Discussion

Insulin resistance, often found with obesity, necessitates the hyperproduction and processing of proinsulin into mature insulin in order to augment insulin secretory capacity, causing significant mitochondrial oxidative and ER stress [8-10]. Insulin resistance also causes a host of physiological changes—including, but not limited to, mild hyperglycemia, hyperinsulinemia, hyperglucagonemia, dyslipidemia, and systemic inflammation. All of these factors impact the beta-cell, and, depending on whether the exposure is acute or chronic, have differential effects on downstream adaptive vs. terminal biological processes. In a compensating beta-cell, mitochondrial and ER stress are ameliorated by cell signaling pathways such as the adaptive unfolded protein response (UPR) and autophagy, which allow the beta-cell to continue to function at a high level. Beta-cell proliferation is stimulated by cell autonomous and non-autonomous factors, allowing for increased beta-cell mass. So long as the beta-cell is able to continue to adapt, T2D will not occur. Yet, many of the factors linked with beta-cell adaptation in the obese & insulin resistant state(s) have also been proposed as contributing to the development and/or pathophysiology of T2D itself, including in our own previously-published work on PGE_2_ production and EP3 signaling [1]. In this work, we have defined a potential mechanism of beta-cell compensation characterized by islet mRNA expression of PTGS2/COX-2, the rate-limiting enzyme for PGE_2_ production, in islets collected from a panel of non-diabetic human organ donors. In this study, enhanced *PTGS2* expression occurs in the absence of either enhanced islet *PTGER3* expression or strong effects of an EP3 antagonist on the insulin secretion response. Instead, *PTGS2* expression is directly correlated with islet insulin content. In other words, the amount of insulin secreted from islets of non-diabetic human donors in all conditions tested is proportional with their *PTGS2* expression, while insulin secretion as normalized to percent of content is unchanged: donor islets with higher *PTGS2* expression secrete more insulin because they have more to secrete. These results identify PGE_2_ (and/or possibly other PTGS2/COX-2 metabolites) as important factors in the maintenance or augmentation of functional beta-cell mass. Consistent with our previously-published work [2], we also found evidence for signaling mechanisms downstream of the EP3-associated Gα_z_ that may regulate beta-cell mass, albeit in an agonist-independent manner.

COX-1 and COX-2 (officially known as prostaglandin-endoperoxidase 1 and 2: PTGS1 and PTGS2), catalyze the rate-limiting step in the production of PGE_2_ derived from arachidonic acid (AA) incorporated in plasma membrane phospholipids. High glucose, free fatty acids, and/or pro-inflammatory cytokines have all been shown to upregulate the expression and/or activity of enzymes involved in the PGE_2_ synthetic pathway, including phospholipase A2 (PLA2: which cleaves AA from membrane phospholipids); COX-1 and COX-2 (which convert arachidonic acid to the intermediate, PGH_2_); and PTGES, PTGES2, and PTGES3, which convert PGH_2_ to PGE_2_ [1, 11-16]. Using BMI as a marker of obesity/insulin resistance, we found a weak correlation with *PTGS2* expression (Figure 1B), consistent with results of a previous study[1]. Yet, BMI is a flawed surrogate for both obesity and insulin resistance. While there exists controversy in the literature, recent reports support IL-6 as an “anti-inflammatory” cytokine that promotes beta-cell adaptation. IL-6 is required for the adaptive proliferative response to HFD-induced glucolipotoxicity by paracrine effects on the alpha-cell [17]. IL-6 was also recently identified as mediating beta-cell autophagy—a survival mechanism working in concert with the adaptive UPR—through stimulation of alpha-cell GLP-1 production [18]. IL-6 expression has also been specifically linked with PGE_2_ produced by COX-2 and not COX-1, which is of particular relevance to the beta-cell, as, in contrast to nearly all other cell types, beta-cell COX-2 expression is constitutive. Although COX-2 can be further induced by various stimuli, COX-2 is the major isoform responsible for both endogenous and stimulated beta-cell PGE_2_ production [15, 16, 19, 20]. In the current work, in non-diabetic human organ donors, *PTGS2* expression was strongly correlated with IL-6 mRNA expression (Figure 3K). Coupled with the relationship between *PTGS2* expression and islet insulin content (Figure 5E), this result provides more evidence for increased COX-2 abundance as a critical component of the beta-cell adaptive response. A limitation of the current work is that we did not directly measure islet PGE_2_ secretion or confirm COX-2 protein levels by Western blot. Yet, in previous work with islets from non-diabetic and T2D human organ donors, the difference in relative *PTGS2* expression determined by qPCR (∼6-fold) was well-matched to the fold-change in total PGE_2_ released per islet during an overnight culture period (∼4.5-fold) [11]. Therefore, we believe *PTGS2* expression is an acceptable surrogate marker for islet PGE_2_ production rate.

Beta-cell stress can also be elicited by islet cell non-autonomous mechanisms that can build off of each other in a feed-forward mechanism to promote beta-cell dysfunction and, ultimately, loss of functional beta-cell mass. The systemic inflammation coincident with the insulin resistance of obesity further promotes mitochondrial and ER stress in already stressed beta-cells by the secretion of pro-inflammatory cytokines from insulin-resistant, M1-polarized macrophages resident within the islet itself, ultimately attracting cytotoxic immune cells and causing beta-cell demise [17, 21, 22]. Resident islet macrophages also have critical roles in beta-cell adaptation and failure. Addition of PGE_2_ to macrophages induces low-level IL-6 expression and promotes the polarization of macrophages to the M2 state [21] and prevents pro-inflammatory cytokine secretion by lipopolysaccharide-activated M1 macrophages [23]. In cancer cells, COX-2-mediated PGE_2_ production suppresses the expression of tumor cell chemokines required for the recruitment of cytotoxic immune cells [23]. Taken together, COX-2-mediated PGE_2_ production and IL-6 expression/secretion by both beta-cells and islet immune cells coordinate to elicit multiple beneficial and overlapping effects on beta-cell function and survival, including through other possible mechanisms not discussed here.

PGE_2_ is the most abundant natural ligand for the EP3 receptor (encoded by the *PTGER3* gene). Unless islets are isolated from human organ that are frankly T2D, islet PTGER3 mRNA expression is minimally changed [1, 24]. Islets isolated from non-diabetic human organ donors have little-to-no functional, proliferative, or survival response to either EP3 agonists or antagonists unless islets have been cultured in glucolipotoxic, pro-inflammatory, or other stress conditions [1, 24, 25]. We have previously published or contributed to two studies characterizing *PTGER3* expression in human islet cDNA samples from non-diabetic human organ donors of varying BMI, with seemingly disparate results. In Kimple and colleagues, when islets were binned by obesity status, *PTGER3* expression was significantly correlated with donor BMI [1]. In the current study, *PTGER3* expression was not correlated with the BMI of the donor, whether linearly or by binning islets by obesity status (Supplementary Table 4). A lack of linear correlation of *PTGER3* expression in the current study with the BMI of non-diabetic donors is consistent with the findings of Carboneau and colleagues [24]. In our experience using mouse models of glucose intolerance and T2D, as well as islets isolated from T2D human organ donors as compared to non-diabetic donors, EP3 mRNA abundance changes of 100-fold or more from the non-diabetic state predict islet responsiveness to EP3 agonists and antagonists [1]. As the mean difference in *PTGER3* expression by obesity status in Kimple and colleagues was relatively small (ΔΔC_t_ ≈ 1.5; fold-change ≈ 2.8), it is likely this change is of limited biological relevance, regardless of statistical significance. Support for this concept is the lack of a potentiating effect of L798,106 on GSIS as donors become increasingly obese (Figure 4B and Supplementary Table 6). Yet, as expected, islets with higher *PTGER3* expression tended to have increased GSIS% in stimulatory glucose with and without Ex-4 when L798,106 was included, although this effect was weak and was only brought out by a population-based analysis (Figure 4B and Figure 5B,C).

In the beta-cell, EP3 is specifically coupled to the inhibitory G protein α-subunit, Gα_z_. Gα_z_ is a classical inhibitory Gα_i/o_ subunit, negatively regulating the production of cAMP by adenylate cyclase (AC). Of all the Gα_i/o_ subfamily members, Gα_z_ is the most disparate, both at the primary sequence level and with regards to its biochemical properties. The GTP hydrolysis (i.e., inactivation) rate of Gα_z_ is incredibly slow (t_1/2_ ≈ 10 min) [26, 27]. Therefore, Gα_z_ has the potential to elicit partial tonic inhibition AC-mediated cAMP production: a well-known contributor to beta-cell function, replication and survival. There are 10 distinct AC isoforms, but *in vivo* Gα_z_ activity has only been demonstrated towards AC1 and AC5, with activity towards AC6 (closely related to AC5) being shown *in vitro* [28]. In this work, *ADCY1, ADCY5*, and *ADCY6* expression were all positively correlated with donor BMI and *CCNA1* expression (Figure 1D-F, Figure 2D,E and Tables 3 and 4), suggesting their importance in the beta-cell compensatory response—a theory also supported in the literature [29-33]. In this work, Gα_z_ expression in islets from non-diabetic human organ donors was correlated with decreased islet insulin content (Figure 5D), with no impact on GSIS% (Supplementary Table 6). In a study of wild-type and Gα_z_-null C57BL/6N mice, extended feeding of a 45 kcal% HFD—a strong model of beta-cell compensation for insulin resistance—Gα_z_ loss synergized with HFD feeding to promote increased beta-cell replication, significantly augmenting beta-cell mass, with no effects on GSIS% [2]. These latter results are fully consistent with our findings on the relationship between *GNAZ* expression and islet insulin content and GSIS% in islets isolated from non-diabetic human organ donors. As we did not directly quantify replication in this study, though, we cannot confirm that increased islet insulin content is due to increased beta-cell mass and not increased insulin content of individual beta-cells, nor can we confirm putative changes in beta-cell mass are due to replication and not survival. Furthermore, as the mRNA expression of Gα_z_-sensitive AC isoforms was only quantified in Set 1 of our human islet panel, and Gα_z_ mRNA expression only in Set 2, we cannot directly compare the relationship of *ADCY1, ADCY5, ADCY6*, and *GNAZ* expression with the full panel of markers of beta-cell compensation, nor can we determine the correlation of *ADCY1, ADCY5*, and *ADCY6* expression with islet insulin content or measurements of beta-cell function.

The decreased insulin content (if representative of beta-cell mass) in Gα_z_ overexpressing islets (Figure 7B) would appear as direct support for an effect of Gα_z_ signaling on human beta-cell proliferation. Yet, as the replication rate of isolated human pancreatic islets is quite low, it is more likely these results are due to a negative effect of Gα_z_ on beta-cell survival. In the INS-1 (832/13) and (832/3) rat insulinoma cell lines, adenoviral overexpression of wild-type human Gα_z_ significantly reduced cAMP production and total cell number after a 2-3-day culture, synergizing with streptozotocin or IL-1β to promote beta-cell apoptosis [4]. Furthermore, in models of T1D, islets from Gα_z_ null mice had significantly fewer TUNEL-positive beta-cells, in addition to a significantly enhanced beta-cell proliferation rate and functional response to glucose and Ex4 [4, 5]. Yet, in the context of HFD-feeding, even wild-type islets from HFD-fed mice had few-to-no TUNEL-positive beta-cells, with the effects of Gα_z_ loss being attributed solely to effects on beta-cell replication ([2] and M.E.K., unpublished data). Effects of Gα_z_ activity on proliferation in the beta-cell highly compensating for insulin resistance and survival in the context of terminal diabetes, particularly when PGE_2_/EP3/Gα_z_ signaling is dysfunctionally up-regulated, are not discordant.

While *GNAZ* expression was negatively correlated with human islet insulin content (Figure 5D), *PTGER3* expression was not (Supplementary Table 6), even though gene expression of the rate-limiting enzymes for PGE_2_ production, COX-1 and COX-2, were positively correlated with both donor BMI and *IL6* expression (Figure 1A,B and Figure 3J,K); the latter a surrogate marker for the beta-cell compensatory state. In fact, *PTGER3* expression was positively correlated with *CCNA1* expression (Figure 3A), supporting, if anything, increased beta-cell replication with increased EP3 expression in non-diabetic human donor islets. There are several non-mutually-exclusive explanations for this discrepancy. First, while heterotrimeric G proteins are traditionally thought of as being activated by ligand binding to their associated receptor, receptor-independent mechanisms have been described [34-36], although none specifically for Gα_z_. Second, in all species characterized, the *PTGER3* gene encodes multiple splice variants differing only in their C-terminal tails: critical determinants of G protein coupling, desensitization, and constitutive activity. The mouse *Ptger3* gene encodes three splice variants (EP3α, EPβ, and EP3γ) that have been much more widely characterized than the human splice variants. EP3α and EP3γ have partial to nearly-full constitutive activity, respectively, against G_i/o_ subfamily members[14, 37]. EP3γ expression has been shown to be up-regulated in aged mouse islets, while islets from aged Gα_z_-null mice have enhanced beta-cell proliferation and function, supporting EP3γ/Gα_z_ signaling as contributing to the mild glucose intolerance of aging [2, 24]. The human *PTGER3* gene encodes 12 known mRNA splice variants (with 4 additional variants being computationally mapped), producing 12 proteins with distinct C-terminal sequences. Of the mouse isoforms, both EP3α and EP3γ have human homologs—EP3I (a.k.a. variant 4) and EP3II (a.k.a. variant 5): the most highly-abundant human EP3 splice variants [14, 38]. Due to the complexity of *PTGER3* splicing and overlapping ORFs in the human genome, we quantified only total *PTGER3* expression. Yet, differences in the relative expression of agonist-sensitive vs. constitutively-active human EP3 splice variants in the healthy vs. compensating beta-cell could explain how changes in Gα_z_ expression could modulate beta-cell mass in an agonist-independent manner without corresponding changes in overall *PTGER3* gene expression. Finally, Gα_z_ is unique in that, its GTP-bound state, it can bind to Rap1GAP [39, 40], a negative regulator of the small G protein, Rap1. Rap1 is well-known as a contributor to the cAMP-mediated amplification pathway of GSIS, but has also been well-characterized as an oncogene [41], and previous work from our laboratory has implicated cAMP-mediated Rap1 activity in a non-canonical mechanistic target of rapamycin complex 1 (mTORC1) signaling pathway that promotes beta-cell replication [42]. Interestingly, binding of Gα_z_ to Rap1GAP and adenylate cyclase are mutually-exclusive [40]. Furthermore, cAMP-, insulin- and inflammation-mediated signaling mechanisms that are prominent in the beta-cell highly compensating for insulin resistance all have been shown to result in post-translational modifications of Gα_z_, Rap1GAP, and Rap1 that impact protein activity and effector preference [43-49]. Of note, when Rap1GAP is phosphorylated by PKA, it is ineffective to catalyze the GTP hydrolysis of Rap1 [46]. Our primary working hypotheses is that, unless PGE_2_ production and EP3 expression are dysfunctionally up-regulated in human islets by the conditions of the T2D state (as described in previous work [1] and mimicked in this work by adenoviral overexpression of Gα_z_), PGE_2_ signaling through EP3 is a functional compensatory mechanism to sequester Gα_z_-GTP from negatively inhibiting AC by encouraging binding to a catalytically-inactive Rap1GAP. Confirming this latter “signaling switch” model is our current focus.

In conclusion, we have defined a putative EP3-independent beta-cell compensation mechanism mediated by IL-6 and COX-2 that may enhance functional beta-cell mass in the islet highly compensating for obesity, inflammation and insulin resistance. Furthermore, we have identified Gα_z_ and its downstream AC targets as critical negative and positive regulators, respectively, of the human beta-cell compensatory response, potentially through regulation of beta-cell replication. The major limitation of our study is the primarily correlative nature between comparisons of BMI, gene expression, and biological outcomes. More work will be needed in order to confirm the importance of these signaling pathways in beta-cell compensation, as well as their specific cellular and molecular mechanisms. Yet, our islet studies are well-designed, rigorous, comprehensive, and our results considered in the context of a broad existing body of work, lending credence to the strength and interpretation of our results. Overall, the model(s) proposed in this manuscript are a significant step forward in synthesizing often disparate results describing the mechanisms behind human beta-cell compensation to obesity and insulin resistance, before the progression to overt T2D.

## 5. Acknowledgements

We wish to thank the many present and former members of the Kimple Laboratory who contributed technical assistance or scientific discussion during the course of these experiments. This work was supported by JDRF Grant 17-2011-608 (to M.E.K.), a Type 1 and Translational Pilot Grant Award from the UW Institute for Clinical and Translational Research (to M.E.K.), a Starter Grant in Translational Medicine and Therapeutics from the PhRMA Foundation, ADA Grant 1-16-IBS-212 (to M.E.K.), and NIH Grant R01 DK102598 (to M.E.K.). The funding bodies had no role in any aspect of the work described in this manuscript. This manuscript is the result of work supported with resources and the use of facilities at the William S. Middleton Memorial Veterans Hospital in Madison, WI. The contents do not represent the views of the U.S. Department of Veterans Affairs or the United States Government.

---

**Supplementary Table 1:** Islet Data for Preparations in Set 1 (Received Oct 2010-Feb 2012)

| Unique Identifier | AGE | SEX | BMI | HbA1c | Origin | Islet Isolation Center | Donor History of diabetes? | Assay |
| --- | --- | --- | --- | --- | --- | --- | --- | --- |
| SAMN08786321 | 27 | M | 19 | Not Reported | IIDP | The Scharp-Lacy Research Institute | No | Gene Expression |
| SAMN08933943 | 55 | M | 19.2 | Not Reported | IIDP | Emory University | No | Gene Expression |
| Not Available* | 58 | F | 20.1 | Not Reported | Beta-Pro | Beta-Pro | No | Gene Expression |
| SAMN08933951 | 51 | F | 23.1 | Not Reported | IIDP | Massachusetts General Hospital | No | Gene Expression |
| SAMN08930052 | 61 | F | 23.2 | Not Reported | IIDP | University of Illinois | No | Gene Expression |
| SAMN08786253 | 65 | M | 24.5 | Not Reported | IIDP | The Scharp-Lacy Research Institute | No | Gene Expression |
| SAMN08933947 | 44 | M | 24.7 | Not Reported | IIDP | Massachusetts General Hospital | No | Gene Expression |
| SAMN08786220 | 48 | M | 24.7 | Not Reported | IIDP | Southern California Islet Cell Resource Center | No | Gene Expression |
| SAMN08933959 | 32 | M | 25.9 | Not Reported | IIDP | University of Pittsburgh | No | Gene Expression |
| SAMN08933959 | 32 | M | 25.9 | Not Reported | IIDP | University of Pittsburgh | No | Gene Expression |
| SAMN08933939 | 40 | F | 26 | Not Reported | IIDP | Massachusetts General Hospital | No | Gene Expression |
| SAMN08930051 | 44 | M | 26.3 | Not Reported | IIDP | University of Illinois | No | Gene Expression |
| SAMN08930051 | 44 | M | 26.3 | Not Reported | IIDP | University of Illinois | No | Gene Expression |
| SAMN08933945 | 39 | M | 27.4 | Not Reported | IIDP | Massachusetts General Hospital | No | Gene Expression |
| SAMN08786303 | 20 | M | 28.3 | Not Reported | IIDP | The Scharp-Lacy Research Institute | No | Gene Expression |
| Not Available* | 44 | F | 29.7 | Not Reported | Beta-Pro | Beta-Pro | No | Gene Expression |
| SAMN08786270 | 38 | M | 29.8 | Not Reported | IIDP | The Scharp-Lacy Research Institute | No | Gene Expression |
| SAMN08786342 | 64 | F | 30 | Not Reported | IIDP | The Scharp-Lacy Research Institute | No | Gene Expression |
| SAMN08786201 | 27 | M | 30 | Not Reported | IIDP | University of Miami | No | Gene Expression |
| SAMN08933935 | 29 | M | 30.2 | Not Reported | IIDP | Massachusetts General Hospital | No | Gene Expression |
| SAMN08933935 | 29 | M | 30.2 | Not Reported | IIDP | Massachusetts General Hospital | No | Gene Expression |
| SAMN08785758 | 56 | M | 30.9 | Not Reported | IIDP | The Scharp-Lacy Research Institute | No | Gene Expression |
| SAMN08786288 | 20 | M | 31.3 | Not Reported | IIDP | University of Miami | No | Gene Expression |
| SAMN08786302 | 51 | F | 31.6 | Not Reported | IIDP | University of Pennsylvania | No | Gene Expression |
| SAMN08786302 | 51 | F | 31.6 | Not Reported | IIDP | University of Pennsylvania | No | Gene Expression |
| SAMN08786218 | 17 | M | 32.5 | Not Reported | IIDP | Southern California Islet Cell Resource Center | No | Gene Expression |
| SAMN08933824 | 49 | F | 33.1 | Not Reported | IIDP | University of Minnesota | No | Gene Expression |
| SAMN08786257 | 48 | F | 33.1 | Not Reported | IIDP | University of Miami | No | Gene Expression |
| SAMN08933825 | 48 | M | 34.4 | Not Reported | IIDP | University of Minnesota | No | Gene Expression |
| Not Available* | 37 | F | 34.4 | Not Reported | Beta-Pro | Beta-Pro | No | Gene Expression |
| SAMN08786315 | 28 | M | 34.5 | Not Reported | IIDP | University of Pennsylvania | No | Gene Expression |
| SAMN08786267 | 37 | F | 34.5 | Not Reported | IIDP | Southern California Islet Cell Resource Center | No | Gene Expression |
| SAMN08933940 | 64 | M | 34.8 | Not Reported | IIDP | Massachusetts General Hospital | No | Gene Expression |
| SAMN08933869 | 16 | F | 35.1 | Not Reported | IIDP | Emory University | No | Gene Expression |
| SAMN08786278 | 54 | F | 36 | Not Reported | IIDP | Southern California Islet Cell Resource Center | No | Gene Expression |
| SAMN08786271 | 40 | F | 36.4 | Not Reported | IIDP | The Scharp-Lacy Research Institute | No | Gene Expression |
| SAMN08930084 | 42 | M | 41.3 | Not Reported | IIDP | University of Illinois | No | Gene Expression |
| SAMN08930084 | 42 | M | 41.3 | Not Reported | IIDP | University of Illinois | No | Gene Expression |
| SAMN08786143 | 29 | M | 42 | Not Reported | IIDP | The Scharp-Lacy Research Institute | No | Gene Expression |
| SAMN08786252 | 36 | F | 42.2 | Not Reported | IIDP | Southern California Islet Cell Resource Center | No | Gene Expression |

\*Beta-pro LLC is no longer in operation and Unique Identifiers cannot be obtained

**Supplementary Table 2:** Islet Data for Preparations in Set 2 (Received Mar 2013-May 2015)

| Unique Identifier | AGE | SEX | BMI | HbA1c | Origin | Islet Isolation Center | Donor History of diabetes? | Assay |
| --- | --- | --- | --- | --- | --- | --- | --- | --- |
| SAMN08930526 | 42 | M | 22.8 | Not Reported | IIDP | University of Illinois | No | Gene Expression and GSIS |
| SAMN08775090 | 32 | M | 23.1 | Not Reported | IIDP | The Scharp-Lacy Research Institute | No | Gene Expression |
| SAMN08784299 | 23 | M | 23.9 | Not Reported | IIDP | The Scharp-Lacy Research Institute | No | Gene Expression and GSIS |
| SAMN08930530 | 48 | F | 24.2 | Not Reported | IIDP | University of Illinois | No | Gene Expression |
| SAMN08930578 | 51 | M | 24.7 | Not Reported | IIDP | University of Illinois | No | Gene Expression |
| SAMN08784380 | 50 | M | 25.7 | Not Reported | IIDP | University of Miami | No | Gene Expression and GSIS |
| SAMN08774963 | 39 | F | 25.9 | Not Reported | IIDP | The Scharp-Lacy Research Institute | No | Gene Expression and GSIS |
| SAMN08775094 | 36 | M | 26 | Not Reported | IIDP | The Scharp-Lacy Research Institute | No | Gene Expression and GSIS |
| SAMN08784374 | 24 | F | 26.6 | Not Reported | IIDP | University of Miami | No | Gene Expression |
| SAMN08784309 | 25 | M | 26.6 | Not Reported | IIDP | The Scharp-Lacy Research Institute | No | Gene Expression and GSIS |
| SAMN08783919 | 19 | M | 26.9 | Not Reported | IIDP | University of Wisconsin | No | Gene Expression and GSIS |
| SAMN08783909 | 53 | M | 27.2 | Not Reported | IIDP | The Scharp-Lacy Research Institute | No | Gene Expression and GSIS |
| SAMN08498443 | 61 | F | 27.6 | Not Reported | IIDP | Southern California Islet Cell Resource Center | No | Gene Expression |
| SAMN08783908 | 52 | M | 29.1 | Not Reported | IIDP | The Scharp-Lacy Research Institute | No | Gene Expression and GSIS |
| SAMN08776514 | 48 | F | 29.2 | Not Reported | IIDP | University of Wisconsin | No | Gene Expression and GSIS |
| SAMN08776501 | 46 | M | 29.3 | Not Reported | IIDP | The Scharp-Lacy Research Institute | No | Gene Expression and GSIS |
| SAMN08784318 | 43 | M | 29.6 | Not Reported | IIDP | University of Pennsylvania | No | Gene Expression |
| SAMN08775091 | 38 | F | 30.6 | Not Reported | IIDP | The Scharp-Lacy Research Institute | No | Gene Expression |
| SAMN08784301 | 43 | M | 30.6 | Not Reported | IIDP | The Scharp-Lacy Research Institute | No | Gene Expression and GSIS |
| SAMN08774955 | 48 | M | 30.7 | Not Reported | IIDP | Southern California Islet Cell Resource Center | No | Gene Expression |
| SAMN08776527 | 58 | F | 31.1 | Not Reported | IIDP | University of Wisconsin | No | Gene Expression |
| SAMN08774969 | 59 | F | 31.3 | Not Reported | IIDP | The Scharp-Lacy Research Institute | No | Gene Expression and GSIS |
| SAMN08774961 | 52 | F | 31.4 | Not Reported | IIDP | Southern California Islet Cell Resource Center | No | Gene Expression and GSIS |
| SAMN08930577 | 50 | M | 31.7 | Not Reported | IIDP | University of Illinois | No | Gene Expression |
| SAMN08774814 | 45 | F | 32.9 | Not Reported | IIDP | University of Wisconsin | No | Gene Expression |
| SAMN08774197 | 52 | M | 33.3 | 4.6 | IIDP | University of Wisconsin | No | Gene Expression |
| SAMN08776521 | 36 | M | 33.8 | Not Reported | IIDP | The Scharp-Lacy Research Institute | No | Gene Expression |
| SAMN08784305 | 52 | M | 34.3 | Not Reported | IIDP | The Scharp-Lacy Research Institute | No | Gene Expression and GSIS |
| SAMN08784314 | 36 | F | 34.8 | Not Reported | IIDP | The Scharp-Lacy Research Institute | No | Gene Expression and GSIS |
| SAMN08775085 | 58 | M | 34.8 | Not Reported | IIDP | University of Wisconsin | No | Gene Expression and GSIS |
| SAMN08776506 | 40 | M | 35.4 | Not Reported | IIDP | University of Pennsylvania | No | Gene Expression and GSIS |
| SAMN08783899 | 25 | M | 35.7 | Not Reported | IIDP | University of Wisconsin | No | Gene Expression and GSIS |
| SAMN08775087 | 45 | M | 36.4 | Not Reported | IIDP | University of Wisconsin | No | Gene Expression |
| SAMN08784317 | 21 | M | 37 | Not Reported | IIDP | University of Wisconsin | No | Gene Expression and GSIS |
| SAMN08774896 | 63 | M | 38.6 | Not Reported | IIDP | University of Wisconsin | No | Gene Expression |
| SAMN08783906 | 40 | M | 38.9 | Not Reported | IIDP | University of Wisconsin | No | Gene Expression and GSIS |
| SAMN08783912 | 51 | M | 38.9 | Not Reported | IIDP | University of Wisconsin | No | Gene Expression |
| SAMN08775048 | 32 | F | 39.4 | Not Reported | IIDP | University of Wisconsin | No | Gene Expression and GSIS |
| SAMN08775030 | 30 | M | 43.7 | Not Reported | IIDP | University of Wisconsin | No | Gene Expression |
| SAMN08774465 | 38 | F | 44.7 | 4.6 | IIDP | University of Wisconsin | No | Gene Expression |

**Supplementary Table 3:** Islet Data for Preparations Used in Adenoviral GSIS Assays (Received Feb - Mar 2019)

| Unique Identifier | AGE | SEX | BMI | HbA1c | Origin | Islet Isolation Center | Donor History of diabetes? | Duration of Disease | Glucose-lowering agents at TOD | Assay |
| --- | --- | --- | --- | --- | --- | --- | --- | --- | --- | --- |
| SAMN10977276 | 52 | M | 27.2 | 5.7 | IIDP | Southern California Islet Cell Resource Center | No | N/A | N/A | Adenoviral GSIS |
| SAMN11157311 | 34 | F | 31.7 | 7.3 | IIDP | University of Wisconsin | Yes | NR | NR | Adenoviral GSIS |
| SAMN11155033 | 51 | M | 24 | 4.7 | IIDP | Southern California Islet Cell Resource Center | No | N/A | N/A | Adenoviral GSIS |

**Supplementary Table 4:** Statistical Results of Gene vs. BMI Analyses.

| **PTGER3** | **PTGS1** | **PTGS2** | **PTGES** | **PTGES2** | **PTGES3** | **GNAZ** | **Linear Fit** |
| --- | --- | --- | --- | --- | --- | --- | --- |
| 0.02069 | 0.09541 | 0.1104 | 0.1009 | 0.04482 | -0.01404 | -0.03421 | **Slope** |
| 0.04347 | 0.05445 | 0.05346 | 0.05539 | 0.03777 | 0.03253 | 0.04259 | **Std. Error** |
| -0.06598 to 0.1074 | -0.01376 to 0.2046 | 0.003574 to 0.2172 | -0.009795 to 0.2115 | -0.03070 to 0.1203 | -0.07995 to 0.05188 | -0.1204 to 0.05201 | **95% CI** |
| 0.00318 | 0.05379 | 0.06341 | 0.04925 | 0.02257 | 0.005007 | 0.01669 | **R-square** |
| 0.6356 | 0.0854 | 0.043 | 0.0733 | 0.2399 | 0.6686 | 0.4269 | **P-value** |
| **PTGER3** | **PTGS1** | **PTGS2** | **PTGES** | **PTGES2** | **PTGES3** | **GNAZ** | **Obesity Status** |
| -9.671 | -11.69 | -6.894 | -10.66 | -6.565 | -3.686 | -6.436 | **BMI < 30** |
| -9.632 | -11.38 | -5.716 | -10.22 | -6.102 | -3.613 | -6.952 | **BMI ≥ 30** |
| 0.03961 ± 0.5097 | 0.3111 ± 0.5286 | 1.178 ± 0.6152 | 0.4348 ± 0.6474 | 0.4632 ± 0.4231 | 0.07374 ± 0.3642 | -0.5153 ± 0.468 | **Difference ± SEM** |
| -0.9742 to 1.053 | -0.7493 to 1.371 | -0.05136 to 2.407 | -0.8586 to 1.728 | -0.3829 to 1.309 | -0.6641 to 0.8116 | -1.463 to 0.4321 | **95% CI** |
| 0.9832 | 0.5588 | 0.0601 | 0.5043 | 0.2279 | 0.8406 | 0.2778 | **P-value** |

| **ADCY1** | **ADCY5** | **ADCY6** | **CCNA1** | **GCK** | **PKM** | **IL6** | **Linear Fit** |
| --- | --- | --- | --- | --- | --- | --- | --- |
| 0.2878 | 0.3071 | 0.2972 | 0.1187 | 0.3603 | -0.02718 | 0.2023 | **Slope** |
| 0.1102 | 0.1122 | 0.09992 | 0.1003 | 0.09219 | 0.06148 | 0.0875 | **Std. Error** |
| 0.05633 to 0.5193 | 0.07147 to 0.5428 | 0.08426 to 0.5102 | -0.09384 to 0.3313 | 0.1666 to 0.5539 | -0.1582 to 0.1039 | 0.02514 to 0.3794 | **95% CI** |
| 0.2749 | 0.294 | 0.371 | 0.08058 | 0.459 | 0.01286 | 0.1233 | **R-square** |
| 0.0176 | 0.0135 | 0.0094 | 0.2536 | 0.001 | 0.6647 | 0.0263 | **P-value** |
| **ADCY1** | **ADCY5** | **ADCY6** | **CCNA1** | **GCK** | **PKM** | **IL6** | **Obesity Status** |
| -11.07 | -11.1 | -15.47 | -12.07 | -9.861 | -4.157 | -12.28 | **BMI < 30** |
| -8.703 | -8.568 | -13.64 | -10.82 | -6.683 | -3.938 | -9.795 | **BMI ≥ 30** |
| 2.362 ± 1.248 | 2.536 ± 1.155 | 1.836 ± 0.9865 | 1.245 ± 0.9537 | 3.178 ± 1.102 | 0.2189 ± 0.5027 | 2.487 ± 0.9526 | **Difference ± SEM** |
| -0.2142 to 4.939 | 0.1527 to 4.92 | -0.2155 to 3.888 | -0.7278 to 3.218 | 0.9048 to 5.452 | -0.8264 to 1.264 | 0.5582 to 4.415 | **95% CI** |
| 0.0706 | 0.038 | 0.0768 | 0.246 | 0.0081 | 0.6677 | 0.0129 | **P-value** |

**Supplementary Table 5:**
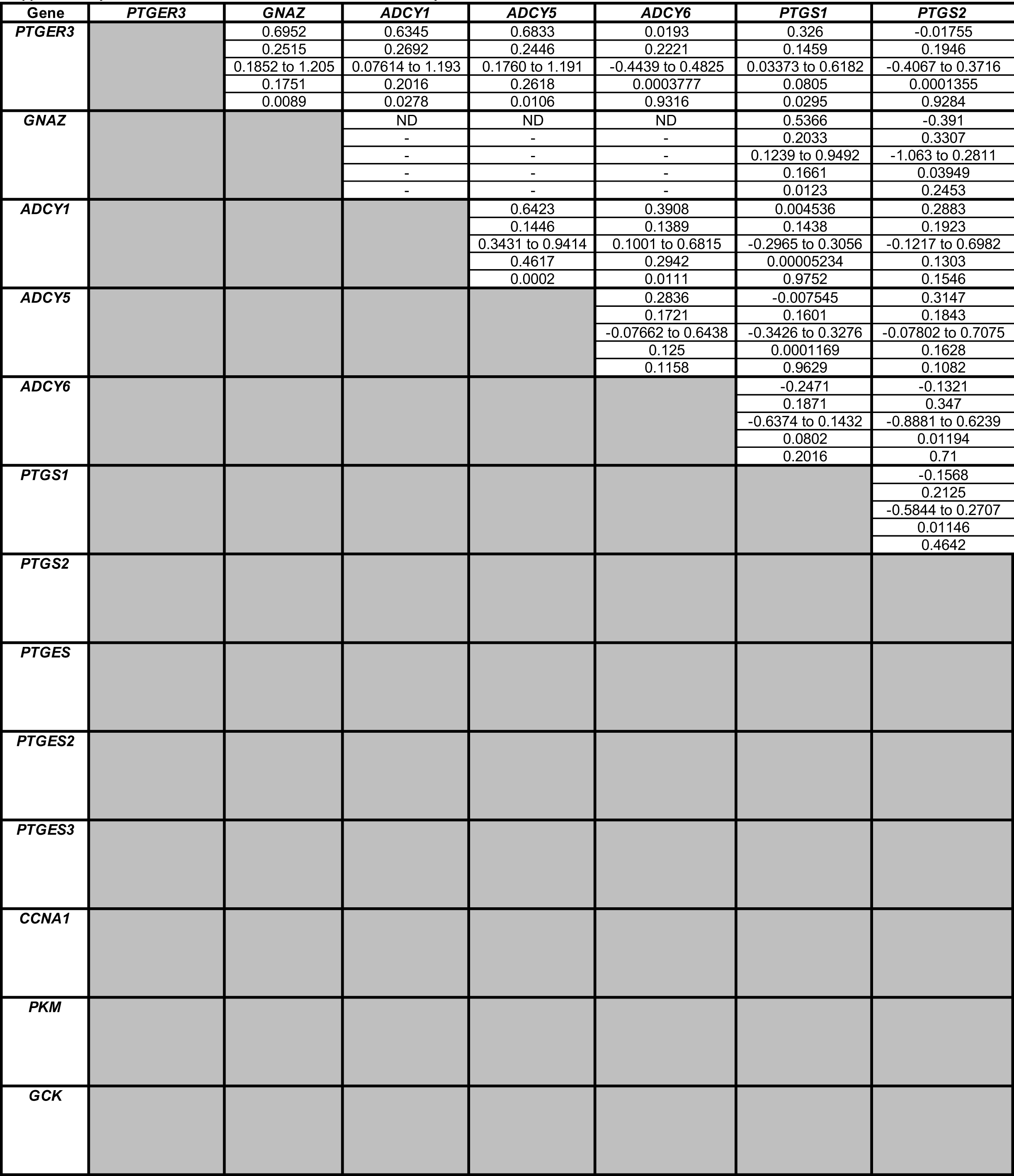
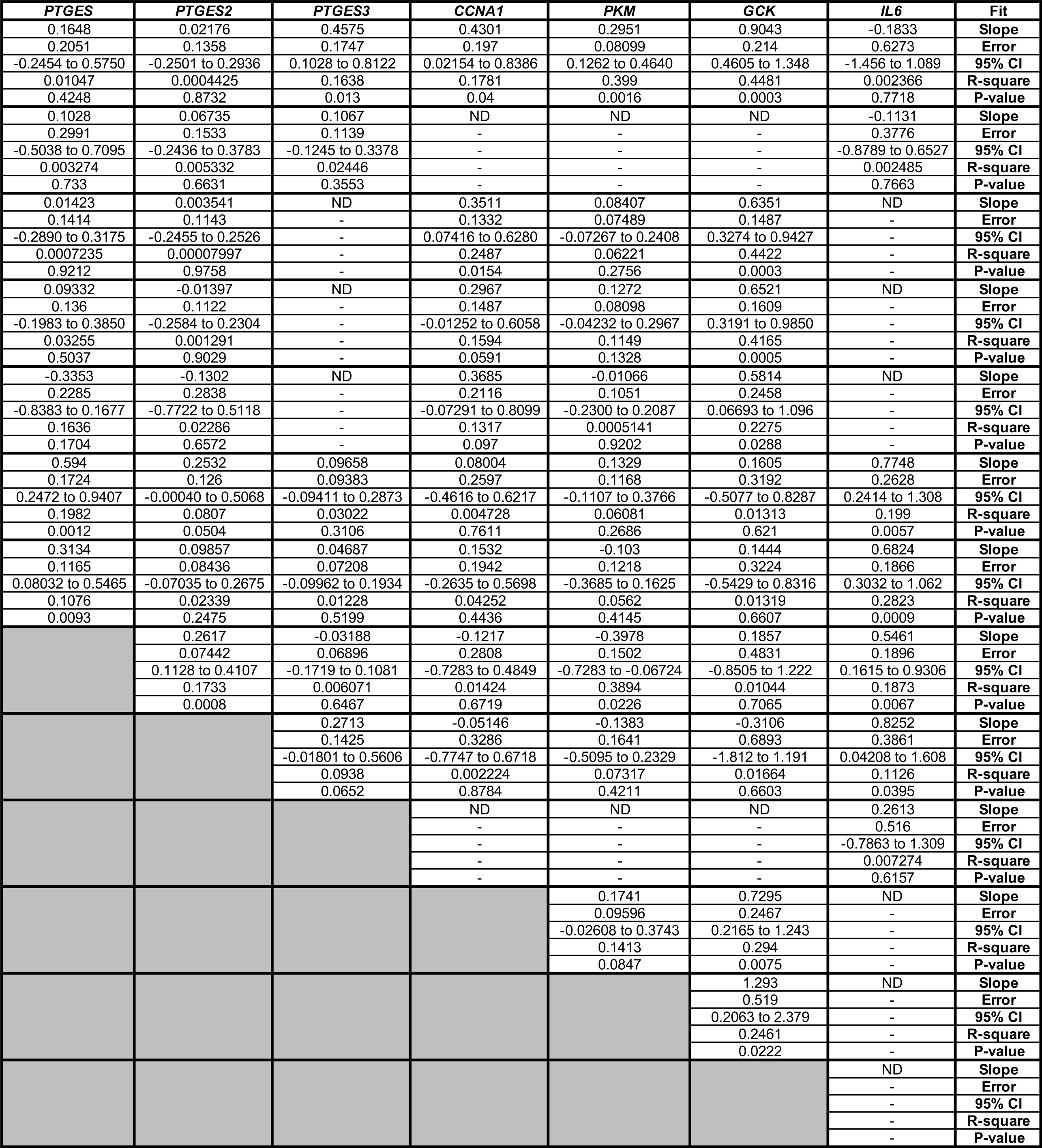
Statistical Results of Gene vs. Gene Anayses.

**Supplementary Table 6:** Statistical Results of BMI and Gene vs. Islet Insulin Content and GSIS Parameters.

| Group | Content<br>(ng/islet) | Total Insulin Secreted (GSIS) (ng/ml) |  |  |  |  |
| --- | --- | --- | --- | --- | --- | --- |
|  |  | 1.7G | 16.7G | 16.7G + Ex4 | 16.7G + L798 | 16.7G+Ex4+L798 |
| **BMI** | 22.36 | 0.2903 | 0.9224 | 0.972 | 1.304 | 1.747 |
|  | 6.942 | 0.1062 | 0.3839 | 0.5001 | 0.4605 | 0.5579 |
|  | 7.879 to 36.84 | 0.06869 to 0.5119 | 0.1216 to 1.723 | -0.07123 to 2.015 | 0.3396 to 2.267 | 0.5794 to 2.915 |
|  | 0.3415 | 0.2718 | 0.224 | 0.1589 | 0.2966 | 0.3404 |
|  | 0.0043 | 0.0128 | 0.0261 | 0.0662 | 0.0107 | 0.0055 |
| **PTGER3** | 6.024 | 0.07021 | 2.355 | 2.169 | 4.036 | 4.694 |
|  | 51.55 | 0.7502 | 2.573 | 3.251 | 3.423 | 4.304 |
|  | -101.5 to 113.5 | -1.495 to 1.635 | -3.012 to 7.723 | -4.612 to 8.950 | -3.129 to 11.20 | -4.314 to 13.70 |
|  | 0.0006824 | 0.0004378 | 0.04021 | 0.02177 | 0.06816 | 0.05892 |
|  | 0.9081 | 0.9264 | 0.3709 | 0.5123 | 0.253 | 0.2891 |
| **GNAZ** | -56.36 | -0.6883 | -2.031 | -2.794 | -1.77 | -2.702 |
|  | 27.53 | 0.4129 | 1.474 | 1.826 | 1.973 | 2.443 |
|  | -113.8 to 1.070 | -1.550 to 0.1729 | -5.106 to 1.044 | -6.603 to 1.016 | -5.900 to 2.359 | -7.814 to 2.411 |
|  | 0.1732 | 0.122 | 0.0867 | 0.1048 | 0.04066 | 0.0605 |
|  | 0.054 | 0.1111 | 0.1835 | 0.1417 | 0.3807 | 0.2825 |
| **PTGS1** | -26.11 | 0.127 | -0.7343 | -0.6927 | 0.1812 | -1.387 |
|  | 40.37 | 0.6052 | 2.116 | 2.673 | 2.854 | 3.559 |
|  | -110.6 to 58.39 | -1.140 to 1.394 | -5.164 to 3.695 | -6.288 to 4.903 | -5.815 to 6.177 | -8.865 to 6.091 |
|  | 0.02155 | 0.002313 | 0.006296 | 0.003521 | 0.0002238 | 0.008367 |
|  | 0.5255 | 0.836 | 0.7324 | 0.7983 | 0.9501 | 0.7013 |
| **PTGS2** | 68.59 | 0.8033 | 2.891 | 3.265 | 3.099 | 4.063 |
|  | 16.32 | 0.272 | 0.9402 | 1.227 | 1.267 | 1.561 |
|  | 34.54 to 102.6 | 0.2360 to 1.371 | 0.9293 to 4.852 | 0.7063 to 5.825 | 0.4464 to 5.752 | 0.7965 to 7.330 |
|  | 0.4689 | 0.3037 | 0.3209 | 0.2616 | 0.2394 | 0.2629 |
|  | 0.0004 | 0.0079 | 0.006 | 0.015 | 0.0244 | 0.0175 |
| **PTGES** | 3.115 | 0.104 | 0.3906 | -0.02051 | -0.3062 | 0.3111 |
|  | 21.19 | 0.3077 | 1.077 | 1.352 | 1.359 | 1.701 |
|  | -41.09 to 47.32 | -0.5378 to 0.7459 | -1.855 to 2.636 | -2.840 to 2.799 | -3.150 to 2.538 | -3.249 to 3.871 |
|  | 0.001079 | 0.005682 | 0.006536 | 0.00001151 | 0.002666 | 0.001758 |
|  | 0.8846 | 0.7388 | 0.7206 | 0.988 | 0.8241 | 0.8568 |
| **PTGES2** | 11.91 | 0.799 | 0.5721 | 0.9 | 1.06 | 0.04261 |
|  | 39.53 | 0.5482 | 2.014 | 2.517 | 2.546 | 3.2 |
|  | -70.55 to 94.36 | -0.3445 to 1.942 | -3.629 to 4.774 | -4.351 to 6.151 | -4.269 to 6.390 | -6.655 to 6.741 |
|  | 0.004515 | 0.09602 | 0.004018 | 0.00635 | 0.009046 | 0.000009331 |
|  | 0.7664 | 0.1605 | 0.7793 | 0.7245 | 0.6817 | 0.9895 |
| **PTGES3** | -29.58 | -0.2525 | -2.426 | -2.219 | -3.089 | -2.702 |
|  | 50.43 | 0.738 | 2.533 | 3.204 | 3.268 | 4.138 |
|  | -134.8 to 75.61 | -1.792 to 1.287 | -7.711 to 2.858 | -8.902 to 4.464 | -9.930 to 3.751 | -11.36 to 5.958 |
|  | 0.01691 | 0.005818 | 0.04385 | 0.02342 | 0.04492 | 0.02195 |
|  | 0.5641 | 0.7358 | 0.3496 | 0.4965 | 0.3564 | 0.5216 |
| **IL6** | 15.93 | 0.2769 | 0.4664 | -0.1233 | -0.1064 | 0.7604 |
|  | 17.48 | 0.2522 | 0.903 | 1.137 | 1.145 | 1.422 |
|  | -20.54 to 52.41 | -0.2492 to 0.8029 | -1.417 to 2.350 | -2.495 to 2.249 | -2.502 to 2.289 | -2.215 to 3.736 |
|  | 0.03987 | 0.05684 | 0.01316 | 0.0005873 | 0.0004547 | 0.01483 |
|  | 0.373 | 0.2853 | 0.6112 | 0.9147 | 0.9269 | 0.599 |

| Insulin Secreted as % of Content (GSIS%) |  |  |  |  |
| --- | --- | --- | --- | --- |
| 1.7G | 16.7G | 16.7G + Ex4 | 16.7G + L798 | 16.7G+Ex4+L798 |
| 0.03574 | 0.0498 | 0.0007139 | 0.1274 | 0.1415 |
| 0.01965 | 0.07774 | 0.1024 | 0.09258 | 0.134 |
| -0.005246 to 0.07673 | -0.1124 to 0.2120 | -0.2129 to 0.2144 | -0.06636 to 0.3212 | -0.1389 to 0.4219 |
| 0.1419 | 0.02011 | 0.00000243 | 0.09065 | 0.05546 |
| 0.0839 | 0.529 | 0.9945 | 0.1848 | 0.3041 |
| -0.03758 | 0.7503 | 0.5273 | 1.111 | 1.567 |
| 0.1276 | 0.4426 | 0.6059 | 0.5728 | 0.8143 |
| -0.3037 to 0.2285 | -0.1730 to 1.674 | -0.7366 to 1.791 | -0.08753 to 2.310 | -0.1371 to 3.272 |
| 0.00432 | 0.1256 | 0.03648 | 0.1654 | 0.1632 |
| 0.7714 | 0.1056 | 0.3945 | 0.0673 | 0.0694 |
| -0.09262 | 0.08495 | 0.05239 | 0.2319 | 0.4302 |
| 0.07217 | 0.2773 | 0.3623 | 0.3521 | 0.4959 |
| -0.2432 to 0.05792 | -0.4935 to 0.6634 | -0.7033 to 0.8081 | -0.5051 to 0.9690 | -0.6077 to 1.468 |
| 0.07609 | 0.004671 | 0.001045 | 0.02232 | 0.0381 |
| 0.214 | 0.7625 | 0.8865 | 0.5181 | 0.3965 |
| 0.02554 | -0.09471 | -0.02766 | 0.02673 | -0.3197 |
| 0.1039 | 0.3836 | 0.4758 | 0.4902 | 0.685 |
| -0.1919 to 0.2430 | -0.8975 to 0.7081 | -1.023 to 0.9682 | -1.003 to 1.057 | -1.759 to 1.119 |
| 0.00317 | 0.003199 | 0.0001779 | 0.0001651 | 0.01196 |
| 0.8085 | 0.8076 | 0.9542 | 0.9571 | 0.6463 |
| 0.06643 | 0.104 | -0.00006566 | 0.07229 | -0.1027 |
| 0.05352 | 0.2043 | 0.2681 | 0.2564 | 0.364 |
| -0.04520 to 0.1781 | -0.3221 to 0.5301 | -0.5594 to 0.5592 | -0.4644 to 0.6089 | -0.8646 to 0.6591 |
| 0.07153 | 0.01279 | 2.998E-09 | 0.004167 | 0.004176 |
| 0.2289 | 0.6162 | 0.9998 | 0.781 | 0.7808 |
| 0.0144 | 0.09516 | -0.09534 | -0.1161 | 0.1111 |
| 0.05248 | 0.1935 | 0.253 | 0.2391 | 0.3406 |
| -0.09508 to 0.1239 | -0.3084 to 0.4988 | -0.6230 to 0.4323 | -0.6165 to 0.3844 | -0.6017 to 0.8240 |
| 0.00375 | 0.01195 | 0.007053 | 0.01225 | 0.00557 |
| 0.7866 | 0.6282 | 0.7102 | 0.6329 | 0.7478 |
| 0.1176 | 0.2409 | 0.2611 | 0.2617 | 0.2814 |
| 0.09465 | 0.3597 | 0.4707 | 0.4483 | 0.6388 |
| -0.07979 to 0.3151 | -0.5094 to 0.9911 | -0.7207 to 1.243 | -0.6765 to 1.200 | -1.056 to 1.618 |
| 0.07171 | 0.02193 | 0.01515 | 0.01762 | 0.01011 |
| 0.2283 | 0.5107 | 0.5852 | 0.5663 | 0.6645 |
| -0.01127 | -0.009999 | 0.07653 | 0.04517 | 0.6162 |
| 0.1261 | 0.4669 | 0.6086 | 0.5912 | 0.8274 |
| -0.2743 to 0.2517 | -0.9838 to 0.9638 | -1.193 to 1.346 | -1.192 to 1.283 | -1.116 to 2.348 |
| 0.0003996 | 0.00002294 | 0.00079 | 0.0003072 | 0.02836 |
| 0.9296 | 0.9831 | 0.9012 | 0.9399 | 0.4656 |
| 0.03045 | 0.02939 | -0.1993 | -0.1133 | 0.007662 |
| 0.04372 | 0.1637 | 0.2089 | 0.2008 | 0.2874 |
| -0.06075 to 0.1216 | -0.3120 to 0.3708 | -0.6351 to 0.2365 | -0.5335 to 0.3069 | -0.5938 to 0.6092 |
| 0.02368 | 0.001609 | 0.04354 | 0.01649 | 0.00003741 |
| 0.4941 | 0.8593 | 0.3514 | 0.579 | 0.979 |

| Fold Change vs. Baseline |  |  |  | Goodness of Fit |
| --- | --- | --- | --- | --- |
| 16.7/1.7 | Ex4/16.7 | L798/16.7 | Ex4+L798/Ex4 |  |
| -0.1144 | -0.02694 | 0.02068 | 0.02479 | Slope |
| 0.2592 | 0.01667 | 0.01425 | 0.01903 | Std Error |
| -0.6569 to 0.4280 | -0.06172 to 0.007845 | -0.009133 to 0.05050 | -0.01504 to 0.06462 | 95% CI |
| 0.01016 | 0.1154 | 0.09987 | 0.08197 | R square |
| 0.6638 | 0.1219 | 0.1628 | 0.2083 | P value |
| 1.731 | -0.1003 | 0.111 | 0.1581 | Slope |
| 1.49 | 0.1045 | 0.09358 | 0.123 | Std Error |
| -1.388 to 4.849 | -0.3183 to 0.1176 | -0.08490 to 0.3068 | -0.09941 to 0.4157 | 95% CI |
| 0.0663 | 0.0441 | 0.0689 | 0.07997 | R square |
| 0.2598 | 0.3483 | 0.2503 | 0.2142 | P value |
| 0.1468 | 0.01064 | 0.04351 | 0.06006 | Slope |
| 0.9395 | 0.0627 | 0.05417 | 0.07155 | Std Error |
| -1.820 to 2.113 | -0.1202 to 0.1414 | -0.06987 to 0.1569 | -0.08970 to 0.2098 | 95% CI |
| 0.001283 | 0.001437 | 0.03283 | 0.03576 | R square |
| 0.8775 | 0.867 | 0.4318 | 0.4117 | P value |
| -0.7473 | 0.01269 | 0.03117 | -0.00816 | Slope |
| 1.261 | 0.08256 | 0.07541 | 0.1033 | Std Error |
| -3.396 to 1.902 | -0.1601 to 0.1855 | -0.1273 to 0.1896 | -0.2251 to 0.2088 | 95% CI |
| 0.01914 | 0.001242 | 0.009403 | 0.0003467 | R square |
| 0.5608 | 0.8795 | 0.6842 | 0.9379 | P value |
| 0.2026 | -0.03226 | -0.002396 | -0.05754 | Slope |
| 0.6718 | 0.04585 | 0.03973 | 0.05088 | Std Error |
| -1.203 to 1.609 | -0.1279 to 0.06338 | -0.08556 to 0.08077 | -0.1640 to 0.04896 | 95% CI |
| 0.004765 | 0.02416 | 0.0001914 | 0.06306 | R square |
| 0.7662 | 0.4898 | 0.9525 | 0.2722 | P value |
| 0.1319 | -0.0469 | -0.0596 | 0.07015 | Slope |
| 0.6337 | 0.04267 | 0.0346 | 0.04651 | Std Error |
| -1.194 to 1.458 | -0.1359 to 0.04212 | -0.1320 to 0.01283 | -0.02720 to 0.1675 | 95% CI |
| 0.002276 | 0.05695 | 0.1351 | 0.1069 | R square |
| 0.8373 | 0.2848 | 0.1012 | 0.148 | P value |
| -0.6967 | -0.01575 | -0.02681 | -0.03719 | Slope |
| 1.199 | 0.08202 | 0.06968 | 0.09213 | Std Error |
| -3.206 to 1.812 | -0.1868 to 0.1554 | -0.1726 to 0.1190 | -0.2300 to 0.1556 | 95% CI |
| 0.01747 | 0.001839 | 0.007729 | 0.008504 | R square |
| 0.5679 | 0.8497 | 0.7047 | 0.6909 | P value |
| -0.3593 | 0.0718 | -0.0005156 | 0.06125 | Slope |
| 1.748 | 0.1042 | 0.09145 | 0.1201 | Std Error |
| -4.018 to 3.299 | -0.1455 to 0.2891 | -0.1919 to 0.1909 | -0.1902 to 0.3127 | 95% CI |
| 0.002219 | 0.0232 | 0.000001673 | 0.01349 | R square |
| 0.8393 | 0.4986 | 0.9956 | 0.6161 | P value |
| 0.2292 | -0.06431 | -0.04393 | 0.0294 | Slope |
| 2.883 | 0.03407 | 0.02964 | 0.04086 | Std Error |
| -5.784 to 6.243 | -0.1354 to 0.006754 | -0.1060 to 0.01811 | -0.05613 to 0.1149 | 95% CI |
| 0.0003159 | 0.1512 | 0.1036 | 0.02652 | R square |
| 0.9374 | 0.0737 | 0.1547 | 0.4807 | P value |

**Supplemental Figure 1.**
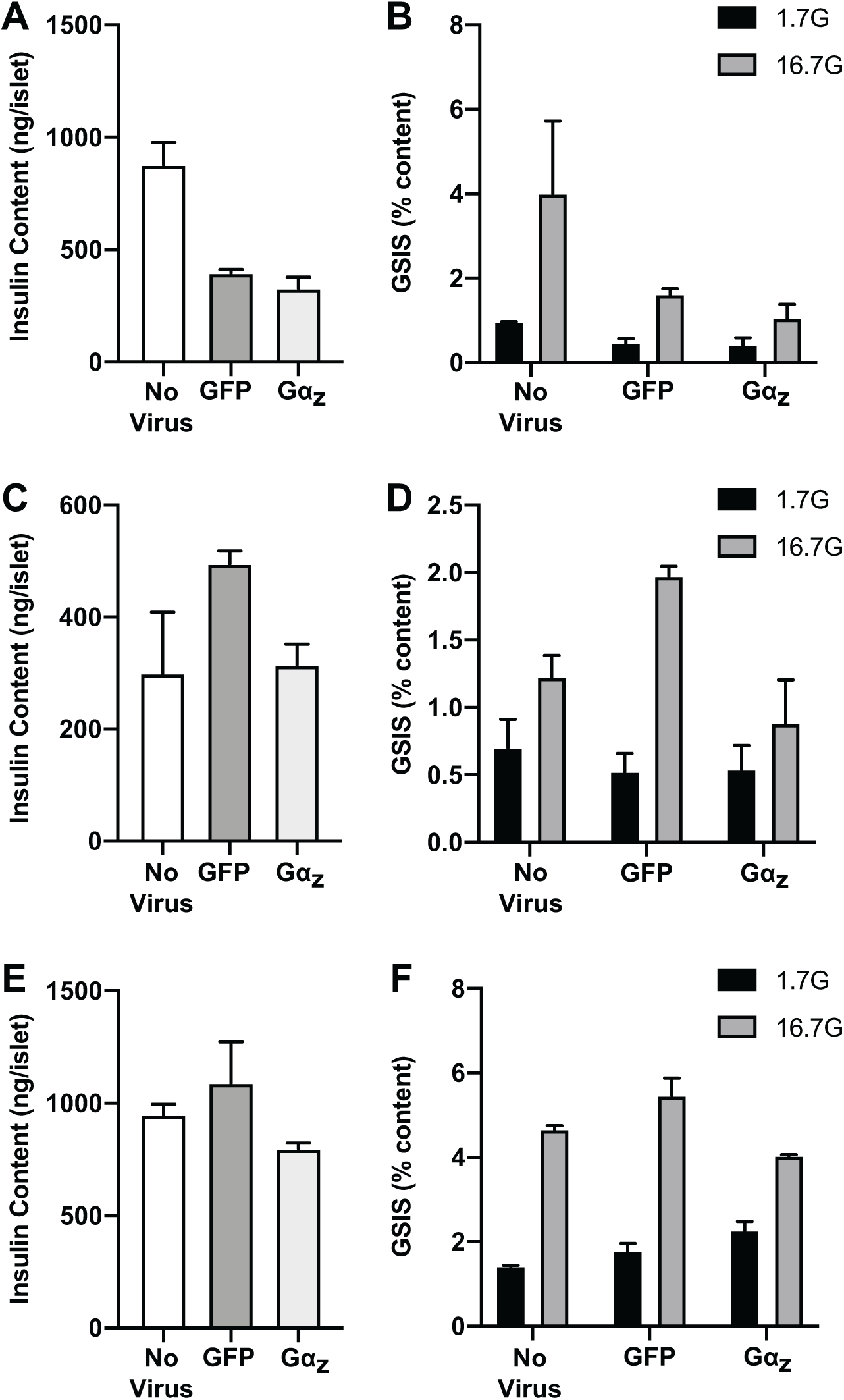
Raw values for islet insulin content and GSIS% for three human islet preparations used in Figure 7, either uninfected or transduced with GFP- or human Gα_z_-encoding adenoviruses. A, C, and E: Total islet insulin content for SAMN10977276, SAMN11157311, and SAMN11155033, respectively. B, D, and F: GSIS as a percent of content in 1.7 mM glucose and 16.7 mM glucose for SAMN10977276, SAMN11157311, and SAMN11155033, respectively. Islet data for each donor’s unique identifier are listed in Supplementary Table 3.

